# Cytosine methylation by ScmA contributes to the fitness of Caulobacter crescentus cells naturally expressing a Vsr-like protein

**DOI:** 10.1101/2025.03.21.644545

**Authors:** Noémie Matthey, Giorgia Wennubst, Karolina Bojkowska, Julien Marquis, Justine Collier

**Affiliations:** Department of Fundamental Microbiology, Faculty of Biology and Medicine, University of Lausanne, Lausanne, CH-1015, Switzerland; Lausanne Genomic Technologies Facility, Faculty of Biology and Medicine, University of Lausanne, Lausanne, CH-1015, Switzerland

## Abstract

While methylated cytosines are known to play important roles in eukaryotes, their significance in bacteria remains poorly understood especially when they are added on genomes by DNA methyltransferases that are not parts of restriction-modification systems. The newly named ScmA protein of *Caulobacter crescentus* is one of these <u>s</u>olitary <u>c</u>ytosine <u>m</u>ethyltransferases. Here, we show that it methylates YGCCGGCR motifs introducing thousands of m5C bases on the *C. crescentus* genome. We also find that wild-type *C. crescentus* cells easily outcompete *ΔscmA* cells during competition experiments. Transcriptome comparisons and using single-cell fluorescent reporters reveal that a DNA damage response is turned on in a significant proportion of *ΔscmA* cells. We then show that this particular response is strictly dependent on the presence of a Vsr-like protein named VsrA. Eventhough the *vsrA* gene is surprisingly neither genetically linked with the *scmA* gene nor with another cytosine methyltransferase gene, VsrA is predicted to encode an endonuclease preventing m5C-to-T mutations. Fluorescence microscopy experiments then show that GFP-VsrA forms foci relatively frequently in stationary phase cells, which may correspond to active repair sites. We propose that VsrA may accidentally create double strand breaks when detecting mismatched bases on the genome of *ΔscmA* cells, leading to their observed loss of fitness. Thus, the presence of *vsr*-like genes can potentially stabilize genes encoding cytosine methyltransferases in bacterial genomes, providing a potential explanation for the relatively high prevalence of solitary cytosine methyltransferases in bacteria even if some of these do not appear to play an easily detectable regulatory role.

## INTRODUCTION

DNA methylation is a process that is widespread throughout all kingdoms of life. Such targeted chemical modifications of bases in genomes can modulate the binding of specific DNA binding proteins and thereby act as epigenetic signals that can influence gene expression (Jeltsch 2002; Breiling and Lyko 2015; Sanchez-Romero and Casadesus 2020). In eukaryotic genomes, C5-methyl-cytosines (m5C) were shown to be linked with a variety of human diseases including immunodeficiency and cancer (Greenberg and Bourc’his 2019). In bacteria, N6-methyl-adenines (6mA), C5-methyl-cytosines (m5C) and/or N4-methyl-cytosines (4mC) are found in ∼93% of sequenced genomes and such methylated bases can represent up to 4% of all the bases of a bacterial genome (Blow et al. 2016; Beaulaurier et al. 2019). As in eukaryotes, methylated bases can influence gene expression through epigenetic processes in bacteria, but they can also play an important role in recognizing self-DNA during horizontal gene transfer events and in protecting bacteria against phage infections (Sanchez-Romero and Casadesus 2020; Anton and Roberts 2021).

DNA methyltransferases (MTases) are the enzymes that can add methyl groups onto bases located in specific DNA motifs during a post-replicative process (Sanchez-Romero et al. 2015). In bacteria, most of the adenine/cytosine MTases are parts of so-called restriction-modification systems (RMS) (Roberts et al. 2015; Blow et al. 2016). In this case, the MTase gene is either located right next to a cognate restriction endonuclease (RE) gene or both activities are encoded by the same gene. The cognate RE recognizes the same DNA motif as the one methylated by the MTase, but only when it is not methylated (Mayo-Munoz et al. 2023). Non-methylated motifs are often found on incoming DNA during horizontal gene transfer (HGT) events or on phage genomes, which is how RMS can act as efficient phage defense mechanisms and limit HGT via RE-mediated DNA cleavage. Still, a significant proportion of the bacterial DNA MTases do not appear to be parts of RMS as they are not encoded by a gene located next to a predicted RE gene; in this case, they are called “solitary” (or “orphan”) DNA MTases (Seshasayee et al. 2012; Blow et al. 2016).

Bacterial solitary DNA MTases may have evolved from degenerate RMS or may have originated from incomplete transfers of mobile genetic elements (MGEs) as many RMS are found in such elements (Seshasayee et al. 2012; Oliveira et al. 2014; Oliveira et al. 2016; Anton and Roberts 2021). Interestingly, eventhough they can then no longer act as RMS, a subset of these have acquired new roles in a diversity of biological processes such as DNA mismatch repair and/or the regulation of chromosomal replication or gene expression (Casadesus and Low 2006; Adhikari and Curtis 2016; Mouammine and Collier 2018; Beaulaurier et al. 2019; Anton and Roberts 2021; Gao et al. 2023). Bacterial epigenetic processes can, for example, regulate cell cycle progression, developmental pathways or virulence. It is however interesting to note that most of the solitary DNA MTases that have been shown to play a regulatory role in bacteria are adenine MTases generating m6A epigenetic signals.

Despite their frequent occurrence, the exact biological role of solitary cytosine MTases creating m5C bases in bacterial genomes usually remains elusive (Anton and Roberts 2021) except for rare examples from *Gammaproteobacteria* and from Group B *Streptococci* (GBS). First, the Dcm cytosine MTase of *Escherichia coli* has been shown to influence stationary phase gene expression, thereby improving the fitness of cells under these conditions (Kahramanoglou et al. 2012; Militello et al. 2014; Militello et al. 2020). Second, the VchM cytosine MTase of *Vibrio cholerae* has been shown to repress the expression of chaperonin-encoding genes (*groESL-2*), playing a role in the tolerance of this bacterium to aminoglycosides (Carvalho et al. 2021) and maybe also in its capacity to infect its hosts (Chao et al. 2015). A last recent example is the Dcm cytosine MTase of GBS that has a major impact on its transcriptome, especially concerning genes encoding proteins involved in carbohydrate transport and metabolism (Manzer et al. 2023).

The frequent occurrence of m5C bases in bacterial genomes is even more intriguing if we consider that these have been shown to be significantly more mutagenic than non-methylated bases, potentially constituting a threat for genome integrity (Lieb and Bhagwat 1996; Kow 2002). Indeed, m5C bases that get accidentally deaminated into T can generate C-to-T mutations if resulting TG mismatches are not detected and repaired on time before the next round of DNA replication. In addition, the spontaneous deamination of m5C occurs at a higher rate than non-modified C, generating C-to-T transition hot-spots at DNA motifs methylated by cytosine MTases in genomes (Lieb and Rehmat 1997). Thus, it has been proposed that cytosine MTases may have a significant impact on the rate of evolution of bacterial genomes (Banerjee and Chowdhury 2006; Estibariz et al. 2019). Still, some bacteria have developed dedicated repair systems, known as <u>v</u>ery <u>s</u>hort <u>p</u>atch (VSP) repair systems, that can detect and then repair TG mismatches in DNA motifs that were initially methylated by cytosine MTase(s) (Hennecke et al. 1991; Lieb and Bhagwat 1996; Bhagwat and Lieb 2002). In these cases, a so-called Vsr endonuclease detects the TG mismatch in the target motif and then catalyzes a nick of the DNA strand containing the T to initiate its removal by a helicase and its replacement with a C through the resynthesis of a short DNA patch by the DNA Pol I. This bacterial VSP repair process was first discovered in *E. coli* where Vsr^Ec^ detects TG mismatches in CCWGG motifs initially methylated by Dcm (Hennecke et al. 1991; Lieb and Bhagwat 1996). In this case, the *vsr* gene is located downstream of its cognate *dcm* gene and co-transcribed from the same operon (Sohail et al. 1990). Later, several other Vsr proteins from a variety of *Neisseria gonorrhoeae* (Kwiatek et al. 2010; Adamczyk-Poplawska et al. 2018) and *Neisseria meningitidis* strains (Bazlekowa et al. 2017) have been characterized. These bacteria often have more than one cytosine MTase and Vsr proteins, but *vsr* genes still appear to stay genetically linked with (sometimes truncated) cytosine MTase genes (solitary or parts of RMS). *In vitro* experiments showed however, that a few of these Vsr proteins can recognize TG mismatches in relatively degenerate motifs (Kwiatek et al. 2010), suggesting that they may be functionally linked with more than one cytosine MTase even if not genetically linked. In some cases, strains carry cytosine MTase genes but no *vsr*-like genes anywhere on the genome (Roberts et al. 2015).

Despite recent progress in the field of bacterial epigenetics/epigenomics, essentially nothing is known about the biological role and the impact of solitary cytosine MTases in the very diverse class of *Alphaproteobacteria* despite their high prevalence (Roberts et al. 2015; Blow et al. 2016). To address this question, the *Caulobacter crescentus Alphaproteobacterium* model appears a prime choice, especially since an analysis of its methylome at different times of its cell cycle has shown that two different DNA motifs carry m5C bases (Kozdon et al. 2013) and since we have already shown that m6A bases on this genome can play a critical role in cell cycle regulation and survival (Gonzalez and Collier 2013; Gonzalez et al. 2014; Mouammine and Collier 2018). Consistent with the existence of two DNA motifs carrying m5C bases, *Caulobacter crescentus* encodes two predicted cytosine MTases: CCNA_03741 and CCNA_01085. Their genes are not genetically linked with a gene annotated to encode an obvious restriction endonuclease and can thus both be classified as solitary DNA MTases (Kozdon et al. 2013). Here, we dissected the potential influence of the <u>s</u>olitary <u>c</u>ytosine <u>M</u>Tase CCNA_01085, now named ScmA, in *C. crescentus*. We constructed a *ΔscmA* mutant and used it to study the impact of ScmA-dependent methylation on *C. crescentus* phenotypes, on its fitness and on its transcriptome. These analyses revealed that ScmA protects cells from DNA damage that can be induced by the CCNA_02930 Vsr-like protein that is now named VsrA. For this reason, wild-type cells largely outcompete *ΔscmA* cells during standard growth conditions. Interestingly, the *vsrA* gene is not genetically linked with the *scmA* gene despite the functional link that we discovered, revealing an original gene positioning architecture compared to previously characterized systems.

## RESULTS

### ScmA methylates the first cytosine in YG<u>C</u>CGGCR motifs

ScmA (CCNA_01085) was predicted to be a <u>s</u>olitary type II <u>c</u>ytosine <u>M</u>Tase since its gene is neither genetically linked with a predicted RE-encoding gene (Fig. 1A), nor encoding a putative endonuclease activity (like RMS IIG) (Marks et al. 2010; Kozdon et al. 2013; Roberts et al. 2015). Consistent with this, we easily constructed a *ΔscmA* mutant strain, demonstrating that ScmA is not essential to protect the *C. crescentus* genome against a lethal RE activity as expected for MTases that are parts of RMS.

**Figure 1:**
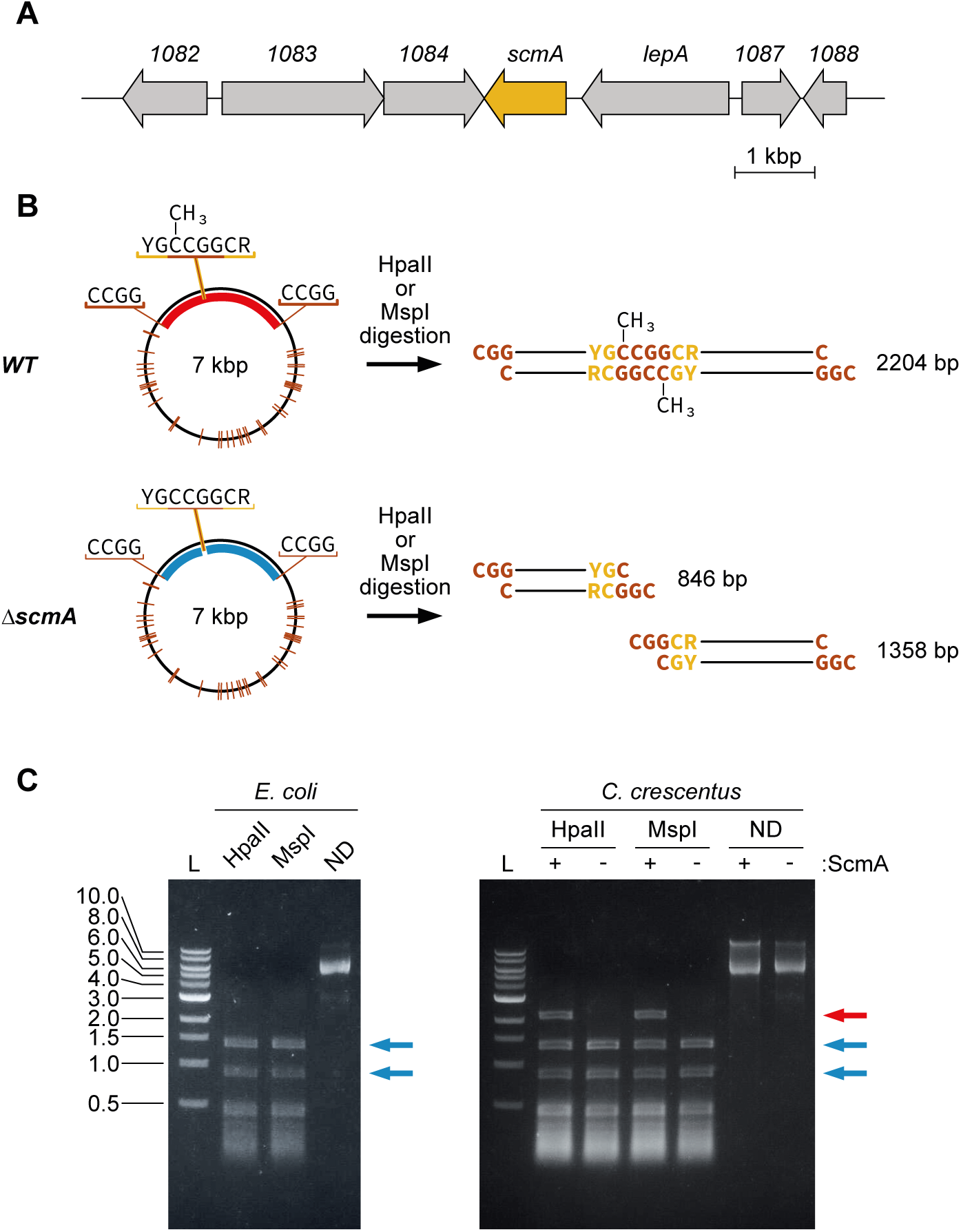
ScmA methylates the first cytosine in YGCCGGCR motifs. **(A)** Schematic showing the organization of genes around *scmA (CCNA_01085)* on the *C. crescentus* NA1000 chromosome. Genes of unknown function are shown using their *CCNA* numbers. (**B**) Map of the pBX-1motif plasmid (left panels) showing the position of the unique YGCCGGCR motif and of the 40 other CCGG motifs (orange lines). These motifs are all cut by the m5C-sensitive HpaII and MspI restriction endonucleases. The right panels show the size (bp) of the expected large restriction fragments following digestion with HpaII or MspI in *WT* or *ΔscmA C. crescentus* cells if ScmA can methylate the first cytosine of the unique YG<u>C</u>CGGCR motif found on that plasmid. (**C**) Image of an agarose gel showing the size of the restriction fragments detected after a digestion of the pBX-1motif plasmid using HpaII, MspI or no enzyme (non-digested; ND). Prior to digestion, pBX-1motif was extracted from *E. coli* TOP10 cells (left panel) or from *WT* (JC450; +) or *ΔscmA* (JC2005; -) *C. crescentus* cells (right panel). Blue arrows highlight the two large restriction fragments (846 and 1358 bp) obtained if the first C in the YGCCGGCR motif is not methylated (HpaII and MspI can then cut the CCGG motif included in that larger motif); the red arrow highlights the unique large restriction fragment obtained (2204 bp) if the first C in the YG<u>C</u>CGGCR motif is methylated (HpaII and MspI can then not cut the <u>C</u>CGG motif included in that larger motif). L: DNA ladder (kbp).

Our previous *C. crescentus* methylome analysis suggested that ScmA may methylate the first C in 5’-YG<u>C</u>CGGCR-3’ motifs since this was the only remaining detected DNA motif containing m5C that is not methylated by the second cytosine MTase (CCNA_03741) found in this bacterium (Kozdon et al. 2013). To confirm this by-default prediction, we used an experimental method based on using m5C-dependent RE to digest DNA with a motif that is expected to contain a modified m5C (Banerjee and Chowdhury 2006). We first constructed a plasmid (named pBX-1motif) containing a single YGCCGGCR motif and introduced it into *E. coli* cells (control) or into *WT* and *ΔscmA C. crescentus* cells. The pBX-1motif plasmid was then extracted from each strain and its capacity to be restricted by HpaII or MspI was tested *in vitro*. HpaII cuts CCGG motifs but only when none of its cytosines are methylated, while MspI cuts non-methylated CCGG motifs (like HpaII) or C<u>C</u>GG motifs with m5C at the second position (Roberts et al. 2023). When the pBX-1motif plasmid was extracted from *E. coli* cells that do not encode a cytosine MTase that can methylate a CCGG motif, the digestion pattern consisted in two large restriction fragments of ∼846 bp and ∼1358 bp (Fig.1C) together with many smaller ones, exactly as expected if neither of the two C found in the unique YGCCGGCR motif engineered on this plasmid are methylated (Fig.1B). Instead, when the pBX-1motif plasmid was extracted from *WT C. crescentus* cells, we observed a third large restriction fragment of ∼2204 bp (=846+1358 bp) when it was digested with either HpaII or MspI (Fig.1C), demonstrating that the first cytosine of the YGCCGGCR located inside this larger restriction fragment was often methylated (m5C), protecting it from digestion by HpaII and MspI (Fig.1B). Importantly, when this plasmid was instead extracted from *ΔscmA C. crescentus* cells, the protected ∼2204 bp fragment could no longer be detected (Fig.1C), demonstrating that ScmA is indeed the cytosine MTase that can methylate the first cytosine of the unique YGCCGGCR motif of this test plasmid in *WT C. crescentus* cells.

It is noteworthy that we could still detect the ∼1358 bp and ∼846 bp restriction fragments, although at lower levels, following the digestion of the test plasmid extracted from *WT C. crescentus* cells (Fig.1C). This observation suggests that a certain proportion (estimated at ∼1/3) of the plasmid molecules were not efficiently methylated by ScmA in the *WT C. crescentus* population from which the plasmid was extracted. This is reminiscent of a previous observation, when we could only detect m5C in ∼76% of the sequencing reads containing YGCCGGCR motifs during SMRT-sequencing analyses after a Tet conversion treatment of the *WT C. crescentus* genome commonly used to detect m5C modifications (Kozdon et al. 2013). Thus, these two independent observations using different methods both suggest that some *C. crescentus* cells may not contain enough ScmA to methylate all the YGCCGGCR motifs found in their genome or that ScmA may not be available to methylate these motifs in a subset of *WT* cells.

Looking more carefully at the *C. crescentus* genome sequence, we found 3054 YGCCGGCR motifs that could be methylated by ScmA, a slightly higher number than expected randomly for the *C. crescentus* genome of ∼4Mbp and ∼67% of G/C content. Still, only 91 (2.98%) of these motifs were mapped in intergenic (IG) regions, demonstrating a significant under-representation since IG regions cover ∼9.6% of the genome.

### Wild-type cells out-compete *ΔscmA cells*

To characterize the impact of DNA methylation by ScmA in *C. crescentus*, we first compared the growth and the morphology of *WT* and *ΔscmA* cells. The growth rates of the two strains cultivated in exponential were very similar in PYE complex medium (Fig.2A, 104 minutes compared to 121 minutes for *WT* and *ΔscmA* cells, respectively) and in M2G minimal medium (Fig.S1, 145 minutes compared to 150 minutes for *WT* and *ΔscmA* cells, respectively). Microscopy analyses of exponentially growing cells cultivated in M2G medium however revealed that *ΔscmA* cells were more often unusually elongated (1.76-fold more frequently with a cell length ≥ 3.5 µm) than *WT* cells (Fig.S2). The cell cycle of these cells may be arrested, or they may be subject to a stress that blocks their division. These elongated cells represented an only limited proportion (6.09%) of the cells in the *ΔscmA* population, suggesting a rather heterogeneous or transient response to the lack of ScmA.

**Figure 2:**
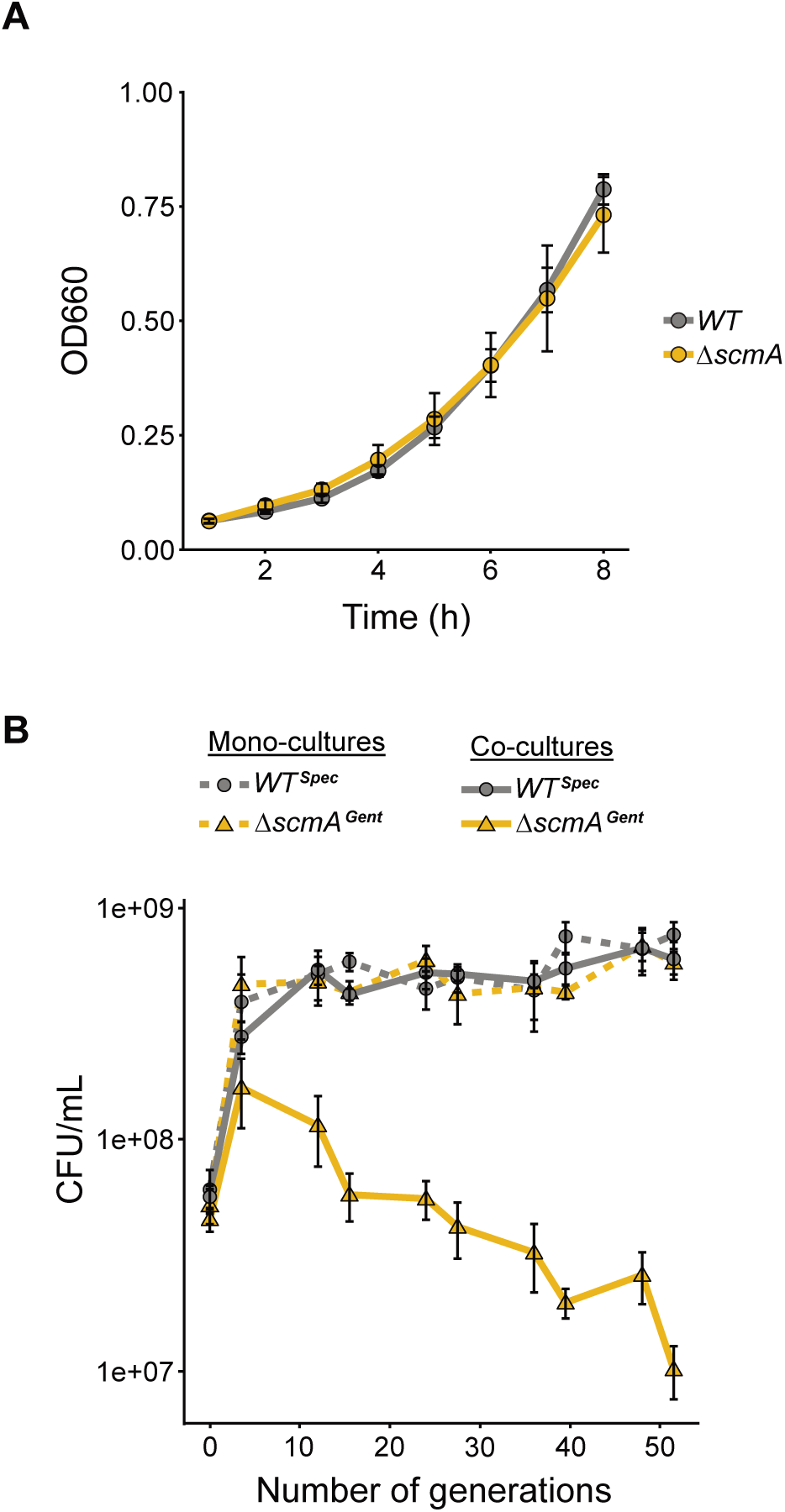
Wild-type *C. crescentus* cells outcompete isogenic *ΔscmA* cells during competition experiments. (**A**) Growth curves of *WT* (JC450) and *ΔscmA* (JC2005) cells cultivated in complex PYE medium. Cells were first cultivated overnight in PYE medium (reaching stationary phase) and cultures were then diluted back into fresh PYE medium to reach an OD_660nm_∼0.05 at time 0. The OD_660_ was then measured every hour for 8 hours. The values plotted in these growth curves correspond to the average measurements from at least three independent biological experiments. (**B**) Monocultures (dashed lines) and co-cultures (solid lines) of *WT^Spec^*(JC2985) and/or Δ*scmA^Gent^* (JC2984) cells were cultivated in complex PYE medium for more than 50 generations (the generation time of these strains is close to 2 hours under these conditions). Before each regular dilution, spectinomycin-resistant (*WT^Spec^*) and gentamycin-resistant (Δ*scmA^Gent^*) colony forming units (CFU) per mL of (co-)cultures were measured on antibiotic-containing PYEA plates. The plotted values correspond to average measurements from four independent biological replicates. In (A) and (B), error bars correspond to standard deviations (± SD).

Considering these very subtle phenotypes, we next decided to compare the fitness of *WT* and *ΔscmA* cells when growing and competing in PYE medium over a prolonged period (∼50 generations). To differentiate the strains from one another, we added spectinomycin or gentamycin resistance cassettes as selective markers in the genomes of *WT* and Δ*scmA* cells, respectively, obtaining the so-called *WT^Spec^* and *ΔscmA^Gent^* strains. To evaluate the relative maintenance of each strain in the mixed population over time, we measured spectinomycin- and gentamycin-resistant colony forming units (CFU)/mL of cultures over time for ∼50 generations making regular dilutions of these co-cultures. Mono-cultures of each strain were also tracked following the same experimental set-up as controls; these confirmed that the antibiotic resistance cassettes added into these strains did not affect their growth (Fig.2B). Strikingly, when the two strains were cultivated together, *WT^Spec^* cells rapidly outcompeted *ΔscmA^Gent^* cells (Fig.2B) demonstrating a strong loss of fitness associated with the loss of ScmA in *C. crescentus*. Indeed, the *ΔscmA^Gent^* cells in the mixed population were rapidly replaced by the *WT^Spec^* cells, with ∼10-fold more *WT^Spec^* cells than *ΔscmA^Gent^* cells after ∼24 generations and ∼60-fold more after ∼50 generations. Thus, *scmA* is not just a cryptic gene that could be easily lost by *C. crescentus* in natural settings when spontaneous mutants/variants usually first compete with their *WT* ancestors before they may find they own favorite niche to propagate.

### Impact of ScmA on the *C. crescentus* transcriptome

Considering that ScmA is a solitary DNA MTase (Fig.1) and that m5C bases can potentially affect gene expression, we first hypothesized that methylation by ScmA may have an impact on the expression of *C. crescentus* genes controlled by promoters that can get methylated by ScmA. As mentioned above, 91 of the YGCCGGCR motifs found on the *C. crescentus* genome are found in IG regions that often contain promoter regions. Thus, to shed light on why *WT* cells can out-compete *ΔscmA* cells (Fig.2B), we decided to compare the transcriptomes of *WT* and *ΔscmA* cells cultivated in exponential phase in M2G medium. RNA-seq experiments revealed that the mRNA levels of 41 genes were significantly different between the two strains (adjusted P value <0.05 and log_2_FC ≤ 1 or ≥ -1 in Fig. 3A and Table S4). Among these, the only gene that was significantly down-regulated in *ΔscmA* cells compared to *WT* cells was *scmA*. Strikingly, when looking at the remaining 40 genes, which were all up-regulated in Δ*scmA* cells compared to *WT* cells, we found an important over-representation (8.4-fold) of the Cluster of Orthologous Group (COG) category L corresponding to “replication, recombination and repair” (Fig.S3). A more detailed analysis (Fig.3A and Table S4) revealed that 52% of these up-regulated genes (21 genes) had been previously shown to be induced in response to DNA damage in *C. crescentus* (da Rocha et al. 2008; Modell et al. 2011; Modell et al. 2014). Examples include the well-conserved *recA* gene (2.2-fold induction in *ΔscmA* compared to *WT* cells), the genes encoding the alternative DNA polymerases *imuA* (6.3-fold induction) and *imuB* (4.6-fold induction) and the gene encoding the apoptosis endonuclease *bapE* (6.6-fold induction). Most of these genes have a LexA-binding box in their promoter region and belong to the SOS-dependent DNA damage response pathway (Galhardo et al. 2005; da Rocha et al. 2008; Modell et al. 2011; Bos et al. 2012; Modell et al. 2014). Still, a few others, including the gene encoding the cell division inhibitor *didA* (5.1-fold induction), belong to other known SOS-independent DNA damage response pathways that are independent of RecA/LexA in *C. crescentus* (Modell et al. 2014). Thus, this transcriptome analysis indicated that DNA damage may take place in at least a subset of *ΔscmA* cells, inducing a general DNA damage response.

**Figure 3:**
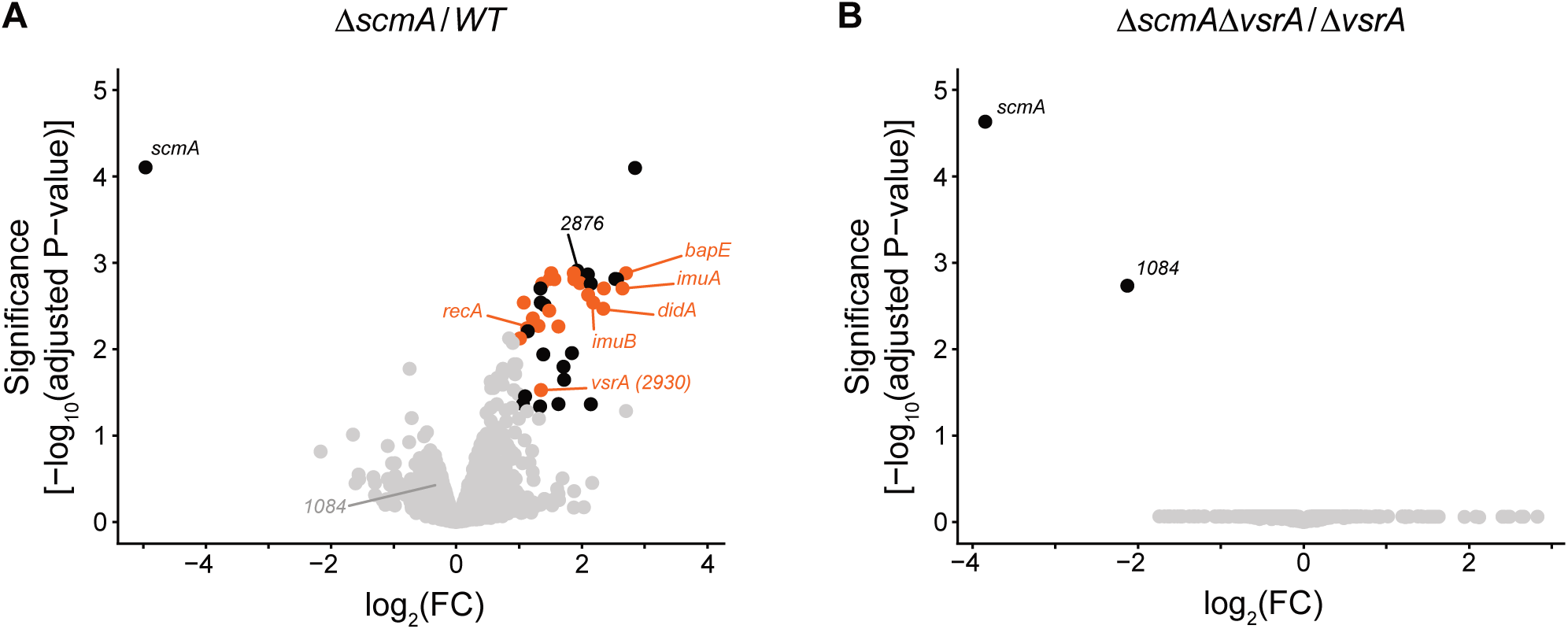
Genes known to be activated in response to DNA damage in *C. crescentus* are activated in Δ*scmA* cells and in a VsrA-dependent manner. Volcano plots showing RNA-Seq results comparing the transcriptomes of *C. crescentus* cells carrying or not the *scmA* gene and cultivated exponentially in M2G medium. FC indicates the fold-change when comparing mRNA levels in cells that do not carry *scmA* compared with cells that do carry *scmA*. Each dot corresponds to one gene. Grey dots correspond to genes that are not considered as differentially regulated (adjusted *P*-value >0.05 and/or -1<log_2_FC<1). Black and red dots correspond to the 41 genes that are differentially regulated (adjusted *P*-value <0.05 and log_2_FC>1 or log_2_FC<-1). Red dots correspond to the 21 differentially regulated genes that are known to be induced during *C. crescentus* DNA damage responses (da Rocha et al. 2008; Modell et al. 2011; Modell et al. 2014). Several non-annotated genes are shown with their *CCNA* numbers. Glimma Volcano plots (interactive HTML graphics) are also available as Supplementary Information. Adjusted *P*-values were calculated based on three independent biological replicates for each strain. **(A)** Volcano plot comparing the transcriptomes of *WT* (JC450) and *ΔscmA* (JC2005) cells. **(B)** Volcano plot comparing the transcriptomes of *ΔvsrA* (JC2540) and *ΔscmA ΔvsrA* (JC2542) cells.

Beyond this COG analysis, we also searched for genes that were significantly up-regulated in *ΔsmcA* cells and that contained at least one YGCCGGCR motif (ScmA target) in their promoter region (250 bp upstream of each open reading frame). Eight genes were found (8/40) (Table S4), including three that also belonged to the LexA regulon (Modell et al. 2014). Thus, altogether, the results of the transcriptome analysis suggested that maximum 5-8 genes may be regulated through the addition of m5C epigenetic marks onto their promoter region by ScmA, or, alternatively, that all the observed mis-regulated genes may simply be linked with the DNA damage response that seemed to be turned on in *ΔscmA* cells.

### An SOS response is turned-on in a subset of *ΔscmA* cells and in a MutL-independent manner

Since we identified many genes known to be activated during the *C. crescentus* SOS response that were also up-regulated in *ΔscmA* cells, we aimed at verifying that an SOS response is indeed turned on in such mutant cells and then also at testing whether this response is homogeneous or heterologous in the *ΔscmA* population.

As a first whole-population reporter of this putative SOS response, we used a plasmid carrying a transcriptional fusion between the *imuA* promoter and the *lacZ* ORF (P*imuA::lacZ*). ImuA is an alternative trans-lesion DNA polymerase that is turned on in a LexA-dependent manner in response to DNA damage in *C. crescentus* (Galhardo et al. 2005). This plasmid was introduced into *WT* and *ΔscmA* cells and the average activity of the P*imuA* promoter in each cell population was measured by β-galactosidase assays using exponentially growing cells cultivated in PYE medium. The results showed that the LexA-dependent P*imuA* promoter was significantly more active in the *ΔscmA* than in the *WT* cell population (1.66-fold induction, Fig.4A), confirming that higher than basal SOS response can be detected in *ΔscmA* cells. However, compared to *WT* cells that were exposed to severe external DNA damaging conditions, such as UV or mitomycin C (Galhardo et al. 2005), this activation remained relatively modest. This observation could be the result of either an SOS response that was turned on at low intensity in all the *ΔscmA* cells, or of a strong SOS response found only in a subset of the *ΔscmA* cells in the population.

**Figure 4:**
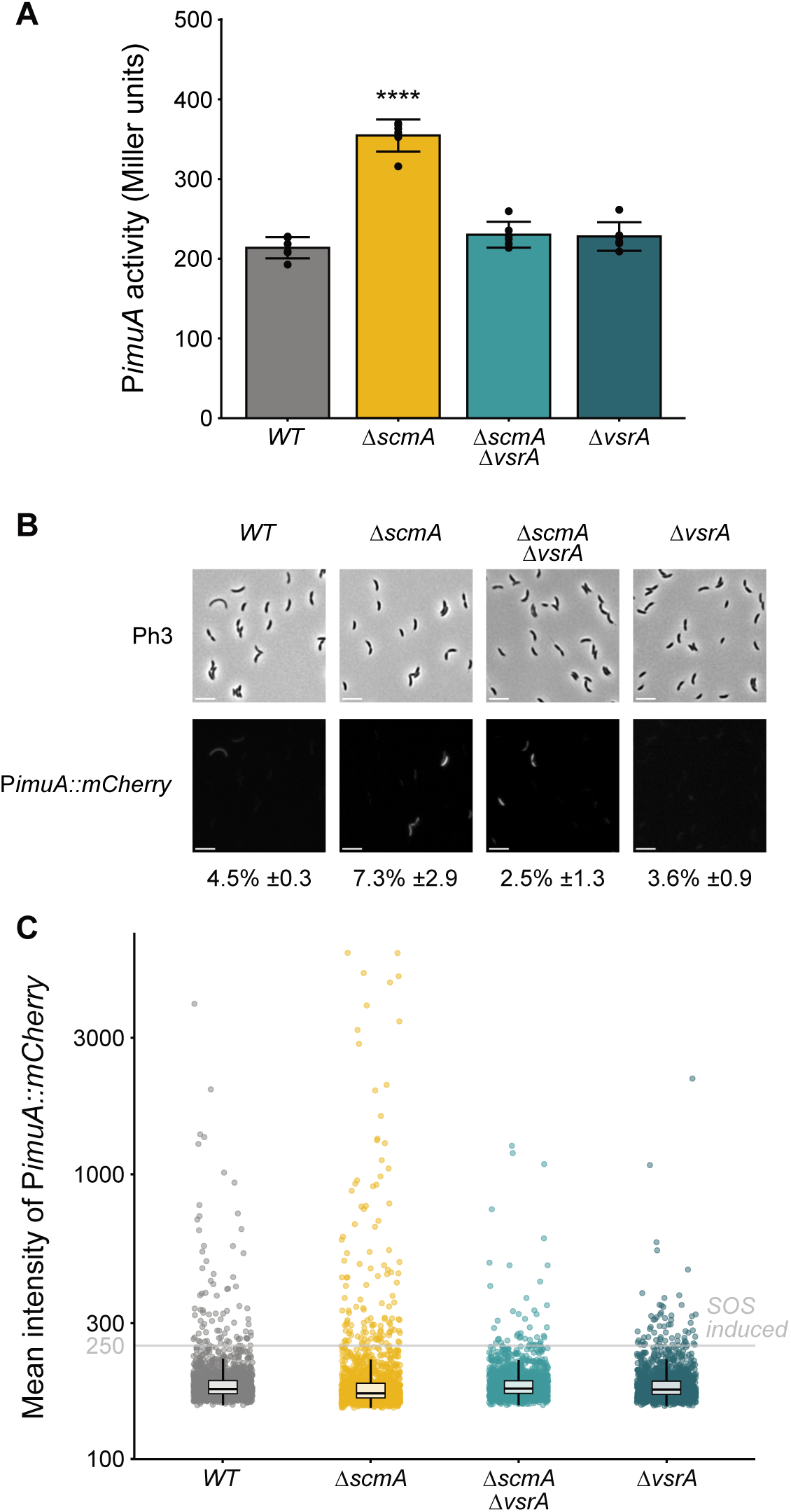
A VsrA-dependent SOS response is turned on in a subset of *ΔscmA* cells. (**A**) β-galactosidase assays showing the VsrA-dependent activation of an SOS response in *ΔscmA* populations of cells. The pP*imuA*::*lacZ*290 plasmid (SOS reporter) was introduced into *WT* (JC450), *ΔscmA* (JC2005), *ΔvsrA* (JC2540) or *ΔscmA ΔvsrA* (JC2542) *C. crescentus* cells. The resulting strains were cultivated in PYE complex medium until cultures reached an 0.25<OD_660nm<_0.3 and β-galactosidase assays were then performed. The plotted promoter activity values (in Miller units) are averages of six independent biological replicates, each with two technical replicates. Error bars correspond to standard deviations (± SD). Statistically significant differences compared to *WT* cells using a student’s *t*-test is indicated as follows: * = *P*-value < 0.05; **** = *P*-value < 0.0001). **(B)** Single-cell fluorescence microscopy assays showing that a VsrA-dependent SOS response is turned on in a subset of *ΔscmA* cells in clonal populations. The P*imuA::mCherry* reporter was integrated at the *imuA* locus on the genome of *WT* (giving JC3104), *ΔscmA* (giving JC3105), *ΔvsrA* (giving JC3107) or Δ*scmA ΔvsrA* (giving JC3109) cells. The resulting strains were cultivated in exponential phase in PYE complex medium. Phase contrast (Ph3) and mCherry images of representative cells are shown. Scale bars indicate 2µm. **(C)** Quantification of the cytoplasmic fluorescence intensity of cells in populations from **(B)**. The average cytoplasmic fluorescence intensity (arbitrary units) of minimum 1600 cells of each strain/culture (4 biological replicates of each with minimum 400 cells/replicate) is shown: the boxes indicate the interquartile range with the center representing the median. Dots above the “SOS” threshold represented by the grey line were used to estimate the percentage of cells in each population displaying an obvious SOS response, as indicated under each microscopy image in panel (B).

To distinguish between these two hypotheses, we constructed a single-cell fluorescent reporter of the SOS response corresponding to a P*imuA::mCherry* construct integrated at the native *imuA* chromosomal locus of *WT* or *ΔscmA* cells. Using this reporter, fluorescence microscopy analyses were used to test whether the SOS response that is turned on in *ΔscmA* cells is homogeneous or heterogeneous in the clonal population. Strikingly, these single-cell assays showed that only a subset of the cells appeared obviously more fluorescent than others in the *ΔscmA* population (Fig.4B&C), indicating that the weak SOS response detected from whole cell populations (Fig.4A) mostly results from a strong SOS induction in a sub-population of cells, rather than from a weak induction in all the cells of the population. This result was further verified using a second single cell fluorescent reporter (P*bapE::mCherry*) of the SOS response (Fig.S4) that gave very similar results.

Interestingly, this observation is reminiscent of what was previously found for *E. coli dam* mutants (Peterson et al. 1985). Dam is also a solitary DNA MTase, but it methylates adenines in GATC motifs on the *E. coli* chromosome. In *E. coli*, m6A bases play an important role during the DNA mismatch repair (MMR) process, when they are used by the MutH endonuclease to distinguish the newly-synthesized DNA strand containing the mismatch that needs to be nicked by MutH during the MMR process (Putnam 2021). In *Δdam* cells, m6A signals are missing and this results in the generation of double-strand breaks (DSB) by MutH that then nicks the two DNA strands instead of one when it tries to repair mismatches from replication errors. Consequently, an SOS response is regularly detected in *Δdam* cells (Bebenek and Janion 1985; Robbins-Manke et al. 2005). Since *C. crescentus* does not have Dam/MutH homologs and since ScmA is also a solitary DNA MTase, we hypothesized that cytosine methylation by ScmA may play a similar role in strand discrimination during the MMR process in *C. crescentus*, potentially explaining why a SOS response is accidentally turned on in a subset of *ΔscmA* cells. If that was the case, one would expect that the SOS response turned on in *ΔscmA* cells would be suppressed in MMR-defective cells. In *C. crescentus*, we recently showed that two proteins contribute to the MMR process: MutS that recognizes the mismatches and MutL that acts as the endonuclease that nicks the newly-synthesized DNA strand containing the replication error to initiate the repair process (Chai et al. 2021) (Fig.S5). To test if MutL may be responsible for the SOS response turned-on in *ΔscmA* cells, we compared the activity of the SOS reporter (P*imuA::lacZ*) in *ΔscmA* and *ΔscmA ΔmutL* mutant cells. β-galactosidase assays showed that the SOS response was essentially identical in both strains (Fig.S6), showing that the MutL endonuclease is not responsible for the SOS response that is turned on in *ΔscmA* cells. Thus, DNA methylation by ScmA most likely does not play a role in the *C. crescentus* MMR process as does Dam in *E. coli*.

### VsrA is responsible for the SOS response turned on in a subset of *ΔscmA* cells

Considering that the MutL endonuclease is not responsible for the SOS response that is turned on in *ΔscmA* cells, we searched for other intracellular sources of potential DNA damage in these cells. Interestingly, our transcriptome analysis revealed that two genes annotated to encode Vsr-like endonucleases were significantly up-regulated in *ΔscmA* cells compared to *WT* cells (Fig.3A and Table S4): *CCNA_02876* (3.8-fold induction) and *CCNA_02930* (2.6-fold induction). To test whether these putative endonucleases may play a role in the SOS response that was observed in *ΔscmA* cells, we deleted one or the other gene in the *WT* and *ΔscmA* strains and then introduced an SOS reporter (P*imuA::lacZ*) into these strains. Strikingly, subsequent β-galactosidase assays revealed that the SOS response was back to basal levels (as *WT*) in *ΔscmA* cells that lacked CCNA_02930 (Fig.4A), while the absence of CCNA_02876 had no impact (Fig.S7). We also confirmed this result using the two fluorescent single-cell reporters P*imuA::mCherry* and P*bapE::mCherry* (Fig.4B, Fig.4C and Fig.S4). Altogether, these results demonstrate that the CCNA_02930 putative endonuclease is fully responsible for the SOS response that is turned on in *ΔscmA* cells.

Looking more carefully at the CCNA_02930 protein sequence (Fig.S8), we found that it belongs to the PD-(D/E)XK superfamily of endonucleases (Steczkiewicz et al. 2012) like the Vsr protein of *E. coli* and that it contains a DUF559 domain, which is often found in Vsr-like endonucleases in a diversity of bacteria. On top of that, a comparison of its predicted 3D structure with that of the well-known *E. coli* Vsr protein (Tsutakawa et al. 1999b) showed that a large part of CCNA_02930 displays similarities. We therefore decided to rename CCNA_02930 VsrA and hypothesized that it may be a VSP repair protein detecting TG mismatches (Fig.5A) like the *E. coli* Vsr protein (Tsutakawa et al. 1999a).

**Figure 5:**
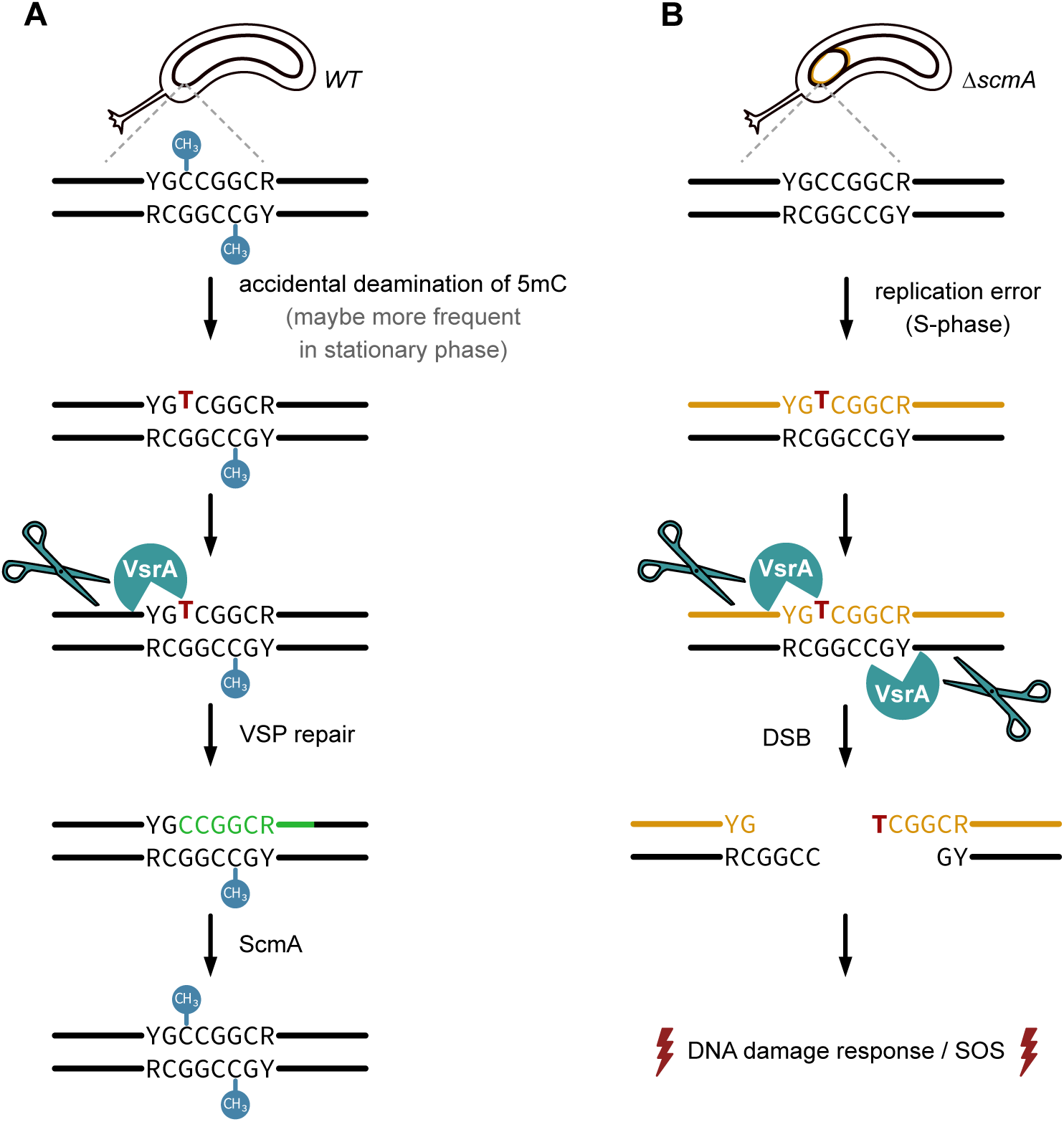
Model for a VsrA-dependent VSP repair process in *C. crescentus*. **(A)** Schematic showing how TG mismatches arising in YGCCGGCR motifs methylated by ScmA (from the accidental m5C deamination into T) may be repaired in VsrA-dependent manner in *WT C. crescentus* cells. The bases replaced during this VSP repair process are shown in green. This model is based on the Vsr-dependent VSP repair mechanism discovered in *E.coli* to repair TG mismatches in DNA motifs methylated by Dcm (Bhagwat and Lieb 2002). **(B)** Schematic showing how VsrA may generate accidental double strand breaks (DSB) if it tries to repair TG mismatches (from DNA replication errors) in non-methylated YGCCGGCR motifs in *ΔscmA C. crescentus* cells. This model is inspired from what was shown to happen when MMR proteins try to repair DNA mismatches in *E. coli Δdam* mutant cells (Bebenek and Janion 1985; Robbins-Manke et al. 2005). The newly synthesized DNA strand is shown in orange in this schematic.

### Cytosine methylation by ScmA does not impact gene expression in the absence of VsrA and no more affects the fitness of *C. crescentus*

Finding that many SOS response genes were turned on in the *ΔscmA* mutant (Fig.3A) complicated our initial goal of determining whether ScmA-dependent methylation may regulate the expression of specific genes in *C. crescentus*. Luckily, observing that the SOS response appeared suppressed in the *ΔscmA ΔvsrA* double mutant (Fig.4) unlocked a novel strategy enabling us to potentially decouple in transcriptomic analyses the VsrA-induced SOS response from the potential direct impact that ScmA methylation may have on the regulation of gene expression. Therefore, to address the initial question, we simply performed RNA-seq experiments comparing the transcriptomes of single *ΔvsrA* and double *ΔscmA ΔvsrA* mutant cells (Fig.3B) in the same growth conditions as before (Fig.3A). Strikingly, our results showed that the only gene that was still significantly mis-regulated (adjusted *P* value <0.05 and log_2_FC ≥ -1) in the *ΔscmA ΔvsrA* strain compared to the *ΔvsrA* strain, beyond the expected *scmA* gene, was the *CCNA_01084* gene (Fig.3B and Table S5), which is located right next to *scmA* on the *C. crescentus* chromosome (Fig.1A). Eventhough this gene is not in the same operon as *scmA* (they are in opposite directions), it is very likely that the deletion of *scmA* that we engineered simply interfered with the normal expression of the nearby *CCNA_01084*. Importantly, its promoter region (250 bp upstream of the ORF) does not include a YGCCGGCR motif (Table S5) ruling out a potential direct impact of ScmA-based methylation on its expression. Taken together, this transcriptome analysis demonstrated that all the genes that were induced in *ΔscmA* cells were linked with the presence of VsrA. Since VsrA is not predicted to be a transcription factor but rather a putative endonuclease (Fig.S8), and since a majority of the genes that were initially induced are known to be activated in response to DNA damage (and without YGCCGGCR motifs in their promoter region) (Fig.3A and Table S4), we hypothesize that VsrA damages the *C. crescentus* chromosome in a subset of *ΔscmA* cells thereby turning on the observed heterogeneous DNA damage response (Fig.4). This important result also indicates that the methylation of cytosines by ScmA on the *C. crescentus* chromosome most likely does not play a role in the regulation of gene expression, at least in the chosen growth conditions.

Considering this information, we hypothesized that the loss of fitness of Δ*scmA* cells compared to *WT* cells (Fig.2B) may then be simply linked to the VsrA-dependent DNA damage response induced in a subset of the mutant cells. To test this, we compared the fitness of Δ*scmA ΔvsrA^Gent^* double mutant cells with that of *WT ^Spec^* cells during competition experiments for over ∼50 generations (Fig.S9). Remarkably, the absence of VsrA completely abolished the deleterious fitness impact linked with the absence of ScmA (Fig.S9).

Altogether, it is thus very likely that the VsrA-dependent SOS response that was turned on in *ΔscmA* cells was responsible for the loss of competitiveness of these mutant cells compared to *WT* cells.

### VsrA forms frequent foci in stationary phase cells

As suggested earlier, the *C. crescentus* VsrA protein may be a VSP repair protein. Considering the functional link that we discovered between ScmA and VsrA and the analogy with the Dcm/Vsr pair in *E. coli*, we now hypothesize that VsrA may recognize TG mismatches arising from the spontaneous deamination of m5C bases in the identified YG<u>C</u>CGGCR motif methylated by ScmA to initiate their repair before they become stable C-to-T mutations during the next round of DNA replication (Fig.5A). In this case, the model in *WT* cells is expected to be that VsrA would nick only the non-methylated DNA strand of the mismatch-containing YGCCGGCR motif containing a T opposite a G instead of a m5C in *WT* cells.

What would then happen in a *ΔscmA* cell if a T is accidentally mis-incorporated instead of a C during DNA replication errors (by the replicative DNA Pol III or by alternative error-prone DNA polymerases) at the third position of a non-methylated YGCCGGCR motif? One can predict that VsrA could end up nicking both DNA strands as neither is methylated in such cells (Fig.5B), exactly as MutH does in *E. coli* Δ*dam* cells during the MMR process (Bebenek and Janion 1985; Robbins-Manke et al. 2005). If this was the case, DSBs would be generated in a subset of the cells in the population (every time a T is accidentally incorporated at the third position of one of the 3054 YG<u>C</u>CGGCR motifs on the *C. crescentus* chromosome) potentially inducing a SOS response to repair them, which is consistent with our findings (Fig.4B&C).

One of our recent studies successfully used fluorescence microscopy to visualize the MMR process in live *C. crescentus* cells (Chai et al, 2021). It showed that a fluorescently tagged MutL endonuclease forms more stable and distinct fluorescent foci when it initiates the repair of mismatches during the MMR process. Following a similar approach, we therefore constructed *C. crescentus* strains expressing a fluorescently tagged VsrA protein (*gfp-vsrA* replacing the native *vsrA* gene) to try to visualize the putative VSP repair process in live *C. crescentus* cells. Fluorescence microscopy analyses showed that GFP-VsrA did form one or more fluorescent foci in ∼20% of the *WT* cells and in ∼6% of the *ΔscmA* cells cultivated exponentially in PYE (Fig.6A&B). These foci appeared at relatively random subcellular positions (Fig.6A) and were not co-localized with the replisome (data not shown), as expected if they were active VSP repair sites located anywhere on the genome. The lower frequency of GFP-VsrA foci in *ΔscmA* compared to *WT* cells may be linked with a lower probability of finding GT mismatches in a YG<u>C</u>CGGCR motif in that strain given that the cytosines are not methylated, although there is not enough evidence to support this hypothesis yet.

**Figure 6:**
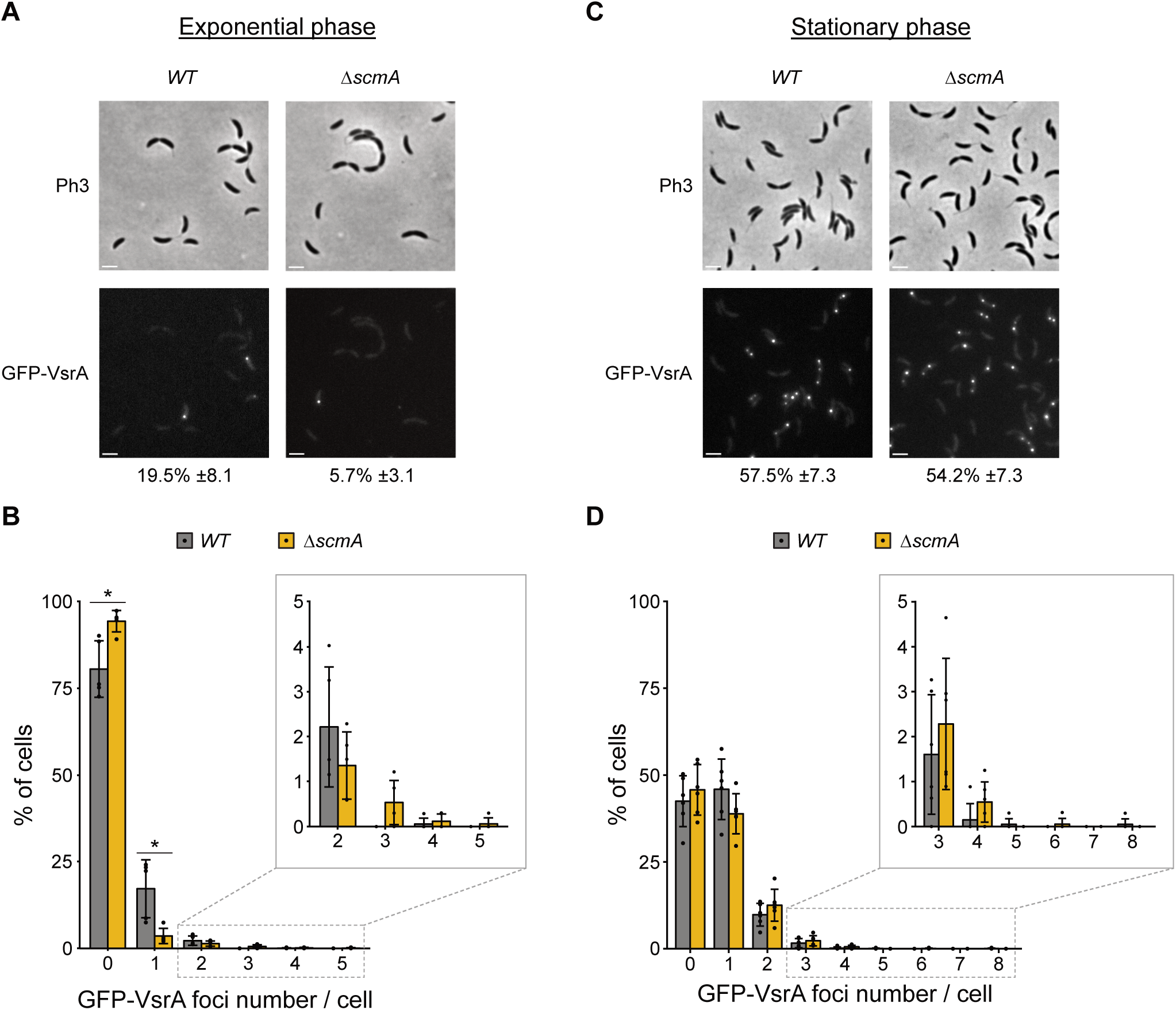
GFP-VsrA forms foci in a majority of stationary phase *C. crescentus* cells. The native *vsrA* gene was replaced by a *gfp-vsrA* construct in *WT* and *ΔscmA* cells, giving the JC2860 and JC2861 strains, respectively. These strains were then cultivated in exponential **(A and B)** or stationary phase **(C and D)** in complex PYE medium before cells were imaged with a fluorescence microscope. Phase contrast (Ph3) and GFP images of representative cells are shown in panels **(A)** and **(C)**. Scale bars correspond to 2µm on these images. The number of detectable GFP-VsrA foci per cell was analyzed for each cell population. Then, the proportion of cells with minimum one detectable GFP-VsrA focus is indicated next to each GFP image. Minimum 300 cells were analyzed for each biological replicate. Panels **(B)** and **(D)** show the percentage of cells displaying a given number of GFP-VsrA foci per cell for each population. The plotted values are averages of at least three independent biological replicates. Error bars correspond to standard deviations (± SD). Statistically significant differences comparing *WT* and *ΔscmA* cells using a student’s *t*-test is indicated as follows: * = *P*-value < 0.05.

Interestingly, previous studies done on *E. coli* cells have shown that the VSP repair process is more active in stationary phase than in exponential phase, either because Vsr levels are higher, or because the accidental conversion of m5C into T may be more frequent during such stress conditions (Macintyre et al. 1999; Bhagwat and Lieb 2002). Supporting this model, we also found that GFP-VsrA formed far more fluorescent foci during stationary than exponential phase in *WT* and *ΔscmA* cells with more than 50% of the cells displaying minimum one focus (Fig.6C&D). If these foci represent active VSP repair sites, it may indicate that TG mismatches occur at higher frequency in *C. crescentus* cells cultivated in stationary phase and/or that VsrA is more abundant/active during stationary phase.

### MMR proteins are most likely not involved in the putative VsrA-dependent VSP repair process

In *E. coli*, but also in *N. gonorrhoeae*, interesting cross-talks have been proposed to take place between the MMR and the VSP repair processes that can, in principle, both detect and repair TG mismatches in DNA motifs carrying m5C (Bhagwat and Lieb 2002; Adamczyk-Poplawska et al. 2018).

First, it was shown that their MutL protein (bridging MutS and MutH in *E. coli*, while acting as the MMR endonuclease in *N. gonorrhoeae*) interacts directly with their Vsr protein(s), potentially impacting the VSP repair process (Mansour et al. 2001; Heinze et al. 2009; Adamczyk-Poplawska et al. 2018). To test whether the *C. crescentus* MMR proteins may interact with the *C. crescentus* VsrA protein, we used bacterial two hybrid assays (BACTH). In brief, we constructed plasmids expressing MutS or MutL proteins fused to the N-terminus of the T18 fragment of the adenylate cyclase and plasmids expressing VsrA fused to the C-terminus of the T25 fragment. These plasmids were then introduced into a BTH101 *E. coli* strain for BACTH assays. Results from these indicate that VsrA can neither interact with MutL, nor with MutS, at least in *E. coli* cells (Fig.S10). Thus, if *C. crescentus* MMR proteins have an influence on the VSP repair process it is probably not through direct interactions between their early actors.

Second, the *E.coli* MutS protein was previously shown to stimulate the VSP repair process *in vivo* and *in vitro,* most likely by helping in the recruitment of Vsr to TG mismatches (Heinze et al. 2009). Conversely, another study indicated that the *N. gonorrhoeae* MutS protein reduces the nicking activity of Vsr near TG mismatches *in vitro*, maybe through a competition mechanism during the recognition of TG mismatches (Adamczyk-Poplawska et al. 2018). Then, to test whether *C. crescentus* MutS protein may affect the capacity of VsrA to potentially cut YG<u>C</u>CGGCR motifs containing TG mismatches on the chromosome of *ΔscmA* cells (Fig.5B), we compared the level of the SOS response in cells expressing or not *scmA* and that lacked MutS (*ΔmutS*) using the P*imuA::lacZ* reporter. We found that the weak SOS response was still induced in response to the loss of ScmA, even in the absence of MutS (1.6-fold induction in the *ΔmutS* background, compared to 1.7-fold induction in the presence of MutS (Fig.S6). Thus, MutS does not seem to promote the VsrA-dependent DNA damage response that takes place in *ΔscmA* cells. Supporting this, we also observed that GFP-VsrA formed fluorescent foci at a relatively similar frequency in cells lacking MutS as in cells expressing *mutS* independently of the presence of the ScmA MTase (Fig.S11A and Fig.S11B) even in cells that were cultivated in stationary phase when these foci are the most visible. Altogether, none of these observations suggest that VsrA may compete or cooperate with MutS in *C. crescentus* cells.

## DISCUSSION

In this study, we used a combination of genetic, genomic and phenotypic analyses to characterize the role of the ScmA solitary cytosine MTase in *C. crescentus*. We used m5C-dependent restriction enzymes to confirm a former prediction (Fig.1) stating that ScmA may methylate the first cytosine in the many YGCCGGCR motifs located on the *C. crescentus* chromosome (Kozdon et al. 2013). A detailed analysis of the phenotypes (Fig.2A and Fig.S1&2) and fitness (Fig.2B) of *ΔscmA* cells revealed that they were significantly less competitive than *WT* cells when co-cultivated in standard growth conditions. First hypothesizing that ScmA-dependent methylation may modulate gene expression through epigenetic mechanisms of regulation like several other solitary MTases (Low and Casadesus 2008; Mouammine and Collier 2018; Sanchez-Romero and Casadesus 2020), we evaluated the impact of ScmA on the *C. crescentus* transcriptome looking for potential explanations for the observed fitness loss. This analysis revealed that a vast majority of the genes that were affected in *ΔscmA* cells encoded proteins that were known to be involved in the adaptation of *C. crescentus* to DNA damage (Fig.3A, Fig.S3 and Table S4). Population-level and single cell reporters were then used to show that an SOS response was indeed turned on (Fig.4A), but only in a subset of *ΔscmA* cells (Fig.4B&C). Among the genes that were significantly activated in *ΔscmA* cells, we identified *vsrA* (Fig.3A and Table S4), encoding a putative Vsr-like endonuclease. Interestingly, this gene was not genetically linked with the *scmA* gene (located at nearly opposite chromosomal positions) unlike the well-characterized *vsr* gene of *E. coli* that is in the same operon as *dcm* (Sohail et al. 1990), and the *vsrA* promoter region (250bp upstream of the ORF) included neither a LexA box, nor a YGCCGGCR motif that can get methylated by ScmA (Table S4). We therefore envisioned that the induction of this putative endonuclease in *ΔscmA* cells may play a role in the apparent DNA damage taking place in a subset of *ΔscmA* cells. Indeed, further transcriptome/single cell SOS reporter assays using double *ΔscmA ΔvsrA* mutant cells clearly demonstrated that the SOS response that is turned on in *ΔscmA* cells is strictly dependent on *vsrA* expression (Fig.3B&4 and Table S5). Furthermore, competition assays demonstrated that the loss of fitness associated with the absence of ScmA was also strictly dependent on *vsrA* expression (Fig.S9). Strikingly, once the *vsrA* gene was removed from *C. crescentus* cells, ScmA had no left-over impact on the expression of genes/operons that appeared to have YGCCGGCR motifs in their promoter region (Table S5). Thus, we can now conclude that the ScmA-dependent methylation of the > 90 promoter/IG regions containing YGCCGGCR motifs on the *C. crescentus* genome does not modulate gene expression, at least in the tested growth conditions.

What is then the biological role of ScmA in *C. crescentus*? One option is that cytosine methylation by ScmA may play an epigenetic regulatory role, but only when *C. crescentus* cells are cultivated in other growth conditions that have not yet been identified. This would be reminiscent of what happens with Dcm in *E. coli* (Militello et al. 2020) and VchM in *V. cholerae* (Chao et al. 2015; Carvalho et al. 2021). ScmA may, alternatively, play a regulatory role in other bacterial strains/species while being only recently acquired by *C. crescentus* from these other bacteria through HGT. Another option is that *scmA* may be a part of a MGE (Oliveira et al. 2014) inducing its own pre-methylation before it moves to other bacterial strains/species by HGT; this would then increase the chances of the MGE to be efficiently transferred into another genome if that genome encodes a RMS that can target the YGCCGGCR motif when it is not methylated. In this case, ScmA would act as a kind of selfish anti-RMS element, promoting the HGT of the MGE. To get insight into this possibility, we used ICEfinder (Liu et al. 2019) and PHASTEST (Wishart et al. 2023) to evaluate whether *scmA* may be included into a prophage or an integrative or conjugative element (ICE) but these tools did not provide evidence that *scmA* may be located within an MGE, even if many genes of unknown function do surround *scmA* on the *C. crescentus* genome (Fig.1A). *scmA* was also not in the prophage identified in the WT/NA1000 *C. crescentus* strain that we used in this study (Marks et al. 2010). It is however still possible that *scmA* may be found in an unknown and relatively original type of MGE that is not yet listed/considered by ICEfinder or PHASTEST. A last option that we believe is worth discussing is that the methylation of cytosines by ScmA may play a role in accelerating the speed of evolution at YGCCGGCR motifs in the genomes of bacteria that encode it, since m5C were previously shown to be C-to-T mutation hot spots in other bacterial species (Lieb and Rehmat 1997; Banerjee and Chowdhury 2006). Although we did not test whether mutations occur at a higher frequency at YGCCGGCR motifs than elsewhere on the *C. crescentus* genome, we have been recently stunned by a study showing that the closely related *C. crescentus* CB15 strain, which is the ancestor of the NA1000 strain that we used in our study (after generations of cultivation in different laboratories around the world), displays only 11 differences compared to NA1000: one consists in a prophage that is integrated into the NA1000 but not into the CB15 genome and the other 10 are single nucleotide polymorphisms (SNPs) (Marks et al. 2010). Strikingly, 2 of these SNPs are C-to-T mutations in YGCCGGCR motifs that are methylated by ScmA in the CB15 and NA1000 strains. Finding that 20% of the SNPs that occurred during this short-term evolution period are specifically located in YGCCGGCR motifs is far more than expected randomly (these motifs represent only 0.6% of the genome). This may just be a coincidence, or it may be seen as first evidence that m5C bases tend to promote the frequency of C-to-T mutations on the *C. crescentus* genome thereby accelerating its evolution despite the presence of a putative VSP repair system.

Is VsrA a VSP repair protein in *C. crescentus*? Finding a functional link between the ScmA-dependent cytosine methyltransferase and the Vsr-like VsrA protein made us hypothesize that VsrA could be a Vsr protein involved in the VSP repair of TG mismatches arising when m5C get accidentally deaminated into T in YG<u>C</u>CGGCR motifs methylated by ScmA (Fig.5A). Supporting the proposition that VsrA can be a functional homolog of the well-characterized Vsr^EC^ protein of *E. coli*, we found significant similarities between the predicted 3D structures of Vsr^Ec^ and VsrA using ChimeraX 1.8 (Fig.S8). In particular, both proteins displayed a four-stranded β-sheet flanked with α-helices on both sides, which is shared by all the members of the PD-(D/E)XK superfamily of nucleases (Steczkiewicz et al. 2012). Based on this predicted structure, the amino-acids located in the nucleolytic active site of Vsr^Ec^ (E25, D51, H64, H69 and D97) could correspond to the E73, D102, D115, H119 and D124 residues of VsrA, suggesting that VsrA may also use magnesium-water clusters to nick DNA motifs as do the three functional Vsr proteins of *E.coli* and *N. gonorrhoeae* do (Tsutakawa et al. 1999a; Tsutakawa et al. 1999b; Kwiatek et al. 2010). The presence of a D115 residue in VsrA instead of the H64 residue in Vsr^Ec^ is reminiscent of what was previously described for the functional Vsr proteins of *N. gonorrhoeae* (V.NgoAXIII) (Kwiatek et al. 2010) and *N. meningitidis* (V.Nme18VIP) (Bazlekowa et al. 2017). Interestingly, *in vitro* experiments showed that these two Vsr proteins can recognize TG mismatches in relatively degenerate DNA motifs that are methylated by any one of the multiple cytosine MTases that are encoded by each of these genomes. Then, we can envision that the *C. crescentus* VsrA may be able to recognize and initiate the repair of TG mismatches located in a variety of DNA contexts. Obviously, more experimental data will be necessary to clarify the exact nucleolytic activity/specificity of VsrA and thereby confirm its role in a VSP repair process in *C. crescentus*. Another interesting outlook of this study will consist in determining whether the GFP-VsrA foci that we distinguished in *C. crescentus* cells (Fig.6) correspond to active VSP repair sites.

Why is VsrA inducing an SOS response in a subset of *ΔscmA* cells? The answer to this question remains elusive but one option is that TG mismatches created by the DNA Pol III during the replication process, and that are not detected on-time by its proofreading activity, can not only be recognized by MutS (Chai et al. 2021), but also by VsrA, at least when located in YGCCGGCR DNA motifs (Fig.5B). In this case, VsrA may cut the two non-methylated DNA strands (Fig.5B), instead of just nicking the only DNA strand that is not methylated as is the case in WT cells (Fig.5A), leading to the formation of accidental DSB and to a transient DNA damage response to promote the repair of this DSB. This hypothesis is reminiscent of what happens in *E. coli dam* mutant cells, where the MutH endonuclease accidentally creates DSB instead of nicking a single DNA strand when trying to repair mismatched bases during the MMR process (Peterson et al. 1985; Robbins-Manke et al. 2005).

A surprising finding during this study was that we discovered a functional link between a cytosine MTase and a Vsr-like protein that are however not encoded by genes that belong to the same operon and that are not even genetically linked on the *C. crescentus* chromosome. This contrasts with the Dcm/Vsr^Ec^ pair found in *E. coli* (Sohail et al. 1990) but, to some extent, shows similarities with a few *Neisseria* Vsr proteins that can repair TG mismatches in a diversity of DNA contexts (Bazlekowa et al. 2017). Still, in *Neisseria*, the genes encoding these characterized Vsr proteins were systematically found next to a MTase gene (that was sometimes truncated), unlike what we found in *C. crescentus*. Such distancing between Vsr- and MTase-encoding genes on bacterial genomes may then promote their horizontal transfer as “solitary” elements and could potentially explain why “solitary” Vsr-encoding genes have now been observed in sequenced bacterial genomes (Roberts et al. 2023). Our study also suggests that the presence of a *vsr*-like gene can stabilize a cytosine MTase-encoding gene in a bacterial genome since we showed that the presence/expression of *vsrA* seriously compromised the fitness/competitiveness of *C. crescentus* cells that lost their *scmA* gene (Fig.2B compared to Fig.S9). Altogether, our study paves the way to better understanding the impact, the transmission and the stabilization of solitary cytosine MTases in a variety of bacteria.

## MATERIALS AND METHODS

### Bacterial growth conditions

Growth conditions are described in Supplementary Information.

### Bacterial strains, plasmids and oligonucleotides

Bacterial strains, plasmids and oligonucleotides used in this study are listed in Tables S1, S2 and S3, respectively, in Supplementary Information.

### Construction of plasmids and strains

Construction of plasmids and strains is described in Supplementary Information.

### Restriction enzyme-based detection of m5C bases

The pBX-1motif plasmid was isolated from overnight cultures of *C. crescentus* cells cultivated in PYE medium or of *E. coli* cells cultivated in LB medium. 700 ng of pBX-1motif were digested for 2h at 37°C using the m5C-sensitive HpaII or MspI restriction endonucleases (New England Biolabs, USA). The size of the resulting restriction fragments was then estimated using agarose gel electrophoresis.

### Fitness assays comparing mono- and co-cultures of *C. crescentus* strains

*C. crescentus* strains were cultivated overnight in PYE medium. Stationary phase mono-cultures were then diluted into 3mL of PYE to reach a starting OD_660nm_ of ∼0.05. For co-cultures to compare the fitness of two strains, an approximately equal numbers of cells of each genotype was inoculated simultaneously into 3mL of PYE to reach a starting OD_660nm_ of ∼0.05 (= OD_660nm_ of ∼0.025 for each strain). Mono- and co-cultures were then diluted back 1000-fold (still in PYE) after 7 hours of growth and again 20-fold after 17 more hours of growth to maintain cells mostly in exponential phase for 24 hours. These 24 hour-cycles were repeated for a total duration of 5 days. The doubling time (dt) of each culture was estimated using the following formula: dt = ln2/r, with r corresponding to the growth rate. We thereby estimated that cells went through ∼51 generations during this 5-day period. At regular time intervals (before each dilution), aliquots of mono- or co-cultures were collected and serially diluted prior to plating onto PYEA+Spec and PYE+Gent to count colony-forming units per mL of culture (CFU/mL) corresponding to each genotype.

### Microscopy and image analysis

Cells were imaged immediately from fresh cultures or after being stored at 4°C for up to a few days after being fixed using a 5X-fix solution (150 mM NaPO4, 12.5% formaldehyde at pH 7.5). Fresh or fixed cells were transferred onto M2 medium with 1% agarose pads on glass slides and were then covered with a coverslip. For phase contrast and fluorescence microscopy, two microscope systems were used depending on the experiments : one (used for Fig.4, Fig.S2 and Fig.S4) was an AxioImager M1 microscope (Zeiss) as described in (Martin et al. 2024) and the other one (used for Fig.6 and Fig.S11) was a Leica DMi8 as described in (Gallay et al. 2021). The Fiji (imageJ) 2.3.0 software with the MicrobeJ plugin (Ducret et al. 2016) was used to analyze images to estimate the length of cells or to identify the proportion of fluorescent cells (from SOS reporters) or of cells with a given number of GFP-VsrA foci in a given cell population.

### RNA extraction

RNA samples were prepared from 10 mL of exponentially growing *C. crescentus* cells cultivated in M2G medium (OD_660nm_ ∼ 0.4). Cells were pelleted and immediately frozen in liquid nitrogen prior to storage at -80°C. RNA were extracted using the RNeasy Mini-Kit from Qiagen following the manufacturer’s protocol including a DNase I (RNase-free DNases set from Qiagen) treatment and a second TURBO DNA-free^TM^ DNase treatment (from Invitrogen) following the protocol of the manufacturer. RNA samples were then re-purified using a RNA clean-up procedure (RNeasy Mini-Kit). Absence of DNA contaminations was verified by standard PCR. The quality of RNA samples was verified on an agarose gel and using a Fragment Analyzer (Agilent Technologies): the RNAs had RQNs between 9.7 and 10.

### RNA-sequencing and analyses

#### RNA-seq library preparation

For Fig.3A and Table S4, RNA-seq libraries were prepared from 1000 ng of total RNA with the Illumina Truseq stranded mRNA Prep reagents (Illumina) using a unique dual indexing strategy and following the official protocol. The polyA selection step was replaced by rRNA depletion step with the Ribo-off rRNA Depletion Kit (Bacteria) (Vazyme). For Fig.3B and Table S5, RNA-seq libraries were prepared from 100 ng of total RNA with the Illumina Stranded mRNA Prep reagents (Illumina) using unique dual indexing strategy, and following the official protocol automated on the Sciclone liquid handling robot (PerkinElmer). The polyA selection step was replaced by rRNA depletion step with the RiboCop for Bacteria mixed bacterial samples reagents (Lexogen). All libraries were quantified by a fluorometric method (QubIT, Life Technologies) and their quality assessed on the Fragment Analyzer. Sequencing was performed either on the Illumina HiSeq 4000 or on the NovaSeq 6000 v1.5 flow cell with single read settings. Sequencing data were demultiplexed using the bcl2fastq2 Conversion Software (version 2.20, Illumina).

#### RNA-seq data processing and analyses

Purity-filtered reads were adapted and quality trimmed with Cutadapt (v. 1.8 for Fig.3A and Table S4 and v. 2.5 for Fig.3B and Table S5) (MARTIN 2011). Upon removal of reads matching to ribosomal RNA sequences (fastq_screen v. 0.11.1) and further filtering for low complexity with reaper (v. 15-065, (Davis et al. 2013), reads were aligned against *Caulobacter crescentus na1000.ASM2200v1* genome using STAR (v. 2.5.3a, (Dobin et al. 2013). The number of read counts per gene locus was summarized with htseq-count (v. 0.9.1, (Anders et al. 2015)) using *Caulobacter crescentus na1000.ASM2200v1* gene annotation. Quality of the RNA-seq data alignment was assessed using RSeQC (v. 2.3.7; (Wang et al. 2012)). The counts per gene table was used for statistical analysis. Quality control analysis was performed through sample density distribution plots, hierarchical clustering and sample PCA. For Fig.3A and Table S4, statistical analyses were performed in R (R version 3.6.1). Genes with no counts were filtered out from the dataset. Library sizes were scaled using TMM normalization (EdgeR package version 3.28.1; (Robinson et al. 2010)) and log2-transformed with the limma cpm function (prior counts set to 1). Differential expression was computed with limma (Limma v3.42.2; (Ritchie et al. 2015)) by fitting the samples into a linear model using as factors three experimental conditions and the batch and then performing the *ΔscmA* vs *WT* comparison. For Fig.3B and Table S5, statistical analyses were performed in R (R version 4.2.1). Genes with at least 1 count per million (cpm) in at least 3 samples were retained for the analysis. Library sizes were scaled using TMM normalization (EdgeR v3.40.1) and log2-transformed with the limma cpm function (prior counts set to 3). Differential expression was computed with limma (Limma v3.54.0) by fitting the samples into a linear model and performing the *ΔscmA ΔvsrA* vs *ΔvsrA* comparison. For both analyses, moderated t-tests were used for the comparisons and P-value adjustments were done using the Benjamini-Hochberg method.

### β-galactosidase assays

*C. crescentus* cells were grown exponentially in the indicated media. 200µl of cultures were used for each β-galactosidase assay following a standard protocol (Miller 1972).

### Statistics and reproducibility

Sample sizes (n) are indicated in figure legends. Statistical analyses were carried out using RStudio (version 2024.04.2+764). Statistical analyses were done using two-tailed Student’s *t*-tests.

### Data availability

Metadata and RNAseq data are available in the following NCBI GEO: GSE286545 (https://www.ncbi.nlm.nih.gov/geo/query/acc.cgi?acc=GSE286545) and GSE286547 (https://www.ncbi.nlm.nih.gov/geo/query/acc.cgi?acc=GSE286547).

## Supporting information

Interactive Volcano Plot Fig.3A

Interactive Volcano Plot Fig.3B

## ACKNOWLEDGMENTS

We thank Tiancong Chai and Jacqueline Masternak for their technical contribution during the construction of strains/plasmids. We also thank Pedro Oliveira, Laurent Casini, Sandra Martin and Florian Fournes for helpful discussions during the project. We finally thank the members of the Lausanne Genomic Technologies Facility for RNA-seq library preparations and sequencing.

## REFERENCES

Adamczyk-Poplawska M, Bandyra K, Kwiatek A. 2018. Activity of Vsr endonucleases encoded by *Neisseria gonorrhoeae* FA1090 is influenced by MutL and MutS proteins. BMC Microbiol 18: 95.

Adhikari S, Curtis PD. 2016. DNA methyltransferases and epigenetic regulation in bacteria. FEMS Microbiol Rev doi:10.1093/femsre/fuw023.

Anders S, Pyl PT, Huber W. 2015. HTSeq--a Python framework to work with high-throughput sequencing data. Bioinformatics 31: 166–169.

Anton BP, Roberts RJ. 2021. Beyond Restriction Modification: Epigenomic Roles of DNA Methylation in Prokaryotes. Annu Rev Microbiol 75: 129–149.

Banerjee S, Chowdhury R. 2006. An orphan DNA (cytosine-5-)-methyltransferase in *Vibrio cholerae*. Microbiology (Reading) 152: 1055–1062.

Bazlekowa M, Adamczyk-Poplawska M, Kwiatek A. 2017. Characterization of Vsr endonucleases from *Neisseria meningitidis*. Microbiology (Reading) 163: 1003–1015.

Beaulaurier J, Schadt EE, Fang G. 2019. Deciphering bacterial epigenomes using modern sequencing technologies. Nat Rev Genet 20: 157–172.

Bebenek K, Janion C. 1985. Ability of base analogs to induce the SOS response: effect of a *dam* mutation and mismatch repair system. Mol Gen Genet 201: 519–524.

Bhagwat AS, Lieb M. 2002. Cooperation and competition in mismatch repair: very short-patch repair and methyl-directed mismatch repair in *Escherichia coli*. Mol Microbiol 44: 1421–1428.

Blow MJ, Clark TA, Daum CG, Deutschbauer AM, Fomenkov A, Fries R, Froula J, Kang DD, Malmstrom RR, Morgan RD et al. 2016. The Epigenomic Landscape of Prokaryotes. PLoS Genet 12: e1005854.

Bos J, Yakhnina AA, Gitai Z. 2012. BapE DNA endonuclease induces an apoptotic-like response to DNA damage in *Caulobacter*. Proc Natl Acad Sci U S A 109: 18096–18101.

Breiling A, Lyko F. 2015. Epigenetic regulatory functions of DNA modifications: 5-methylcytosine and beyond. Epigenetics Chromatin 8: 24.

Carvalho A, Mazel D, Baharoglu Z. 2021. Deficiency in cytosine DNA methylation leads to high chaperonin expression and tolerance to aminoglycosides in *Vibrio cholerae*. PLoS Genet 17: e1009748.

Casadesus J, Low D. 2006. Epigenetic gene regulation in the bacterial world. Microbiol Mol Biol Rev 70: 830–856.

Chai T, Terrettaz C, Collier J. 2021. Spatial coupling between DNA replication and mismatch repair in *Caulobacter crescentus*. Nucleic Acids Res 49: 3308–3321.

Chao MC, Zhu S, Kimura S, Davis BM, Schadt EE, Fang G, Waldor MK. 2015. A Cytosine Methyltransferase Modulates the Cell Envelope Stress Response in the *Cholera* Pathogen [corrected]. PLoS Genet 11: e1005666.

da Rocha RP, Paquola AC, Marques Mdo V, Menck CF, Galhardo RS. 2008. Characterization of the SOS regulon of *Caulobacter crescentus*. J Bacteriol 190: 1209–1218.

Davis MP, van Dongen S, Abreu-Goodger C, Bartonicek N, Enright AJ. 2013. Kraken: a set of tools for quality control and analysis of high-throughput sequence data. Methods 63: 41–49.

Dobin A, Davis CA, Schlesinger F, Drenkow J, Zaleski C, Jha S, Batut P, Chaisson M, Gingeras TR. 2013. STAR: ultrafast universal RNA-seq aligner. Bioinformatics 29: 15–21.

Ducret A, Quardokus EM, Brun YV. 2016. MicrobeJ, a tool for high throughput bacterial cell detection and quantitative analysis. Nat Microbiol 1: 16077.

Estibariz I, Overmann A, Ailloud F, Krebes J, Josenhans C, Suerbaum S. 2019. The core genome m5C methyltransferase JHP1050 (M.Hpy99III) plays an important role in orchestrating gene expression in Helicobacter pylori. Nucleic Acids Res 47: 2336–2348.

Galhardo RS, Rocha RP, Marques MV, Menck CF. 2005. An SOS-regulated operon involved in damage-inducible mutagenesis in *Caulobacter crescentus*. Nucleic Acids Res 33: 2603–2614.

Gallay C, Sanselicio S, Anderson ME, Soh YM, Liu X, Stamsas GA, Pelliciari S, van Raaphorst R, Denereaz J, Kjos M et al. 2021. CcrZ is a pneumococcal spatiotemporal cell cycle regulator that interacts with FtsZ and controls DNA replication by modulating the activity of DnaA. Nat Microbiol 6: 1175–1187.

Gao Q, Lu S, Wang Y, He L, Wang M, Jia R, Chen S, Zhu D, Liu M, Zhao X et al. 2023. Bacterial DNA methyltransferase: A key to the epigenetic world with lessons learned from proteobacteria. Front Microbiol 14: 1129437.

Gonzalez D, Collier J. 2013. DNA methylation by CcrM activates the transcription of two genes required for the division of *Caulobacter crescentus*. Mol Microbiol 88: 203–218.

Gonzalez D, Kozdon JB, McAdams HH, Shapiro L, Collier J. 2014. The functions of DNA methylation by CcrM in *Caulobacter crescentus*: a global approach. Nucleic Acids Res 42: 3720–3735.

Greenberg MVC, Bourc’his D. 2019. The diverse roles of DNA methylation in mammalian development and disease. Nat Rev Mol Cell Biol 20: 590–607.

Heinze RJ, Giron-Monzon L, Solovyova A, Elliot SL, Geisler S, Cupples CG, Connolly BA, Friedhoff P. 2009. Physical and functional interactions between *Escherichia coli* MutL and the Vsr repair endonuclease. Nucleic Acids Res 37: 4453–4463.

Hennecke F, Kolmar H, Brundl K, Fritz HJ. 1991. The vsr gene product of *E. coli* K-12 is a strand- and sequence-specific DNA mismatch endonuclease. Nature 353: 776–778.

Jeltsch A. 2002. Beyond Watson and Crick: DNA methylation and molecular enzymology of DNA methyltransferases. Chembiochem 3: 274–293.

Kahramanoglou C, Prieto AI, Khedkar S, Haase B, Gupta A, Benes V, Fraser GM, Luscombe NM, Seshasayee AS. 2012. Genomics of DNA cytosine methylation in *Escherichia coli* reveals its role in stationary phase transcription. Nat Commun 3: 886.

Kow YW. 2002. Repair of deaminated bases in DNA. Free Radic Biol Med 33: 886–893.

Kozdon JB, Melfi MD, Luong K, Clark TA, Boitano M, Wang S, Zhou B, Gonzalez D, Collier J, Turner SW et al. 2013. Global methylation state at base-pair resolution of the *Caulobacter* genome throughout the cell cycle. Proc Natl Acad Sci U S A doi:10.1073/pnas.1319315110.

Kwiatek A, Luczkiewicz M, Bandyra K, Stein DC, Piekarowicz A. 2010. *Neisseria gonorrhoeae* FA1090 carries genes encoding two classes of Vsr endonucleases. J Bacteriol 192: 3951–3960.

Lieb M, Bhagwat AS. 1996. Very short patch repair: reducing the cost of cytosine methylation. Mol Microbiol 20: 467–473.

Lieb M, Rehmat S. 1997. 5-Methylcytosine is not a mutation hot spot in nondividing *Escherichia coli*. Proc Natl Acad Sci U S A 94: 940–945.

Liu M, Li X, Xie Y, Bi D, Sun J, Li J, Tai C, Deng Z, Ou HY. 2019. ICEberg 2.0: an updated database of bacterial integrative and conjugative elements. Nucleic Acids Res 47: D660–D665.

Low DA, Casadesus J. 2008. Clocks and switches: bacterial gene regulation by DNA adenine methylation. Curr Opin Microbiol 11: 106–112.

Macintyre G, Pitsikas P, Cupples CG. 1999. Growth phase-dependent regulation of Vsr endonuclease may contribute to 5-methylcytosine mutational hot spots in *Escherichia coli*. J Bacteriol 181: 4435–4436.

Mansour CA, Doiron KM, Cupples CG. 2001. Characterization of functional interactions among the *Escherichia coli* mismatch repair proteins using a bacterial two-hybrid assay. Mutat Res 485: 331–338.

Manzer HS, Brunetti T, Doran KS. 2023. Identification of a DNA-cytosine methyltransferase that impacts global transcription to promote group B streptococcal vaginal colonization. mBio 14: e0230623.

Marks ME, Castro-Rojas CM, Teiling C, Du L, Kapatral V, Walunas TL, Crosson S. 2010. The genetic basis of laboratory adaptation in *Caulobacter crescentus*. J Bacteriol 192: 3678–3688.

Martin M. 2011. Cutadapt removes adapter sequences from high-throughput sequencing reads. EMBnetjournal 17: 10–12.

Martin S, Fournes F, Ambrosini G, Iseli C, Bojkowska K, Marquis J, Guex N, Collier J. 2024. DNA methylation by CcrM contributes to genome maintenance in the *Agrobacterium tumefaciens* plant pathogen. Nucleic Acids Res 52: 11519–11535.

Mayo-Munoz D, Pinilla-Redondo R, Birkholz N, Fineran PC. 2023. A host of armor: Prokaryotic immune strategies against mobile genetic elements. Cell Rep 42: 112672.

Militello KT, Finnerty-Haggerty L, Kambhampati O, Huss R, Knapp R. 2020. DNA cytosine methyltransferase enhances viability during prolonged stationary phase in *Escherichia coli*. FEMS Microbiol Lett 367.

Militello KT, Mandarano AH, Varechtchouk O, Simon RD. 2014. Cytosine DNA methylation influences drug resistance in *Escherichia coli* through increased *sugE* expression. FEMS Microbiol Lett 350: 100–106.

Miller JH. 1972. Experiments in Molecular Genetics. Cold Spring Harbor Laboratory Press, Cold Spring Harbor, NY.

Modell JW, Hopkins AC, Laub MT. 2011. A DNA damage checkpoint in *Caulobacter crescentus* inhibits cell division through a direct interaction with FtsW. Genes Dev 25: 1328–1343.

Modell JW, Kambara TK, Perchuk BS, Laub MT. 2014. A DNA damage-induced, SOS-independent checkpoint regulates cell division in *Caulobacter crescentus*. PLoS Biol 12: e1001977.

Mouammine A, Collier J. 2018. The impact of DNA methylation in Alphaproteobacteria. Mol Microbiol doi:10.1111/mmi.14079.

Oliveira PH, Touchon M, Rocha EP. 2014. The interplay of restriction-modification systems with mobile genetic elements and their prokaryotic hosts. Nucleic Acids Res 42: 10618–10631.

Oliveira PH, Touchon M, Rocha EP. 2016. Regulation of genetic flux between bacteria by restriction-modification systems. Proc Natl Acad Sci U S A 113: 5658–5663.

Peterson KR, Wertman KF, Mount DW, Marinus MG. 1985. Viability of *Escherichia coli* K-12 DNA adenine methylase (*dam*) mutants requires increased expression of specific genes in the SOS regulon. Mol Gen Genet 201: 14–19.

Putnam CD. 2021. Strand discrimination in DNA mismatch repair. DNA Repair (Amst) 105: 103161.

Ritchie ME, Phipson B, Wu D, Hu Y, Law CW, Shi W, Smyth GK. 2015. limma powers differential expression analyses for RNA-sequencing and microarray studies. Nucleic Acids Res 43: e47.

Robbins-Manke JL, Zdraveski ZZ, Marinus M, Essigmann JM. 2005. Analysis of global gene expression and double-strand-break formation in DNA adenine methyltransferase- and mismatch repair-deficient *Escherichia coli*. J Bacteriol 187: 7027–7037.

Roberts RJ, Vincze T, Posfai J, Macelis D. 2015. REBASE--a database for DNA restriction and modification: enzymes, genes and genomes. Nucleic Acids Res 43: D298–299.

Roberts RJ, Vincze T, Posfai J, Macelis D. 2023. REBASE: a database for DNA restriction and modification: enzymes, genes and genomes. Nucleic Acids Res 51: D629–D630.

Robinson MD, McCarthy DJ, Smyth GK. 2010. edgeR: a Bioconductor package for differential expression analysis of digital gene expression data. Bioinformatics 26: 139–140.

Sanchez-Romero MA, Casadesus J. 2020. The bacterial epigenome. Nat Rev Microbiol 18: 7–20.

Sanchez-Romero MA, Cota I, Casadesus J. 2015. DNA methylation in bacteria: from the methyl group to the methylome. Curr Opin Microbiol 25: 9–16.

Seshasayee AS, Singh P, Krishna S. 2012. Context-dependent conservation of DNA methyltransferases in bacteria. Nucleic Acids Res 40: 7066–7073.

Sohail A, Lieb M, Dar M, Bhagwat AS. 1990. A gene required for very short patch repair in *Escherichia coli* is adjacent to the DNA cytosine methylase gene. J Bacteriol 172: 4214–4221.

Steczkiewicz K, Muszewska A, Knizewski L, Rychlewski L, Ginalski K. 2012. Sequence, structure and functional diversity of PD-(D/E)XK phosphodiesterase superfamily. Nucleic Acids Res 40: 7016–7045.

Tsutakawa SE, Jingami H, Morikawa K. 1999a. Recognition of a TG mismatch: the crystal structure of very short patch repair endonuclease in complex with a DNA duplex. Cell 99: 615–623.

Tsutakawa SE, Muto T, Kawate T, Jingami H, Kunishima N, Ariyoshi M, Kohda D, Nakagawa M, Morikawa K. 1999b. Crystallographic and functional studies of very short patch repair endonuclease. Mol Cell 3: 621–628.

Wang L, Wang S, Li W. 2012. RSeQC: quality control of RNA-seq experiments. Bioinformatics 28: 2184–2185.

Wishart DS, Han S, Saha S, Oler E, Peters H, Grant JR, Stothard P, Gautam V. 2023. PHASTEST: faster than PHASTER, better than PHAST. Nucleic Acids Res 51: W443–W450.

