## Supplementary Information for "Cytosine methylation by ScmA contributes to the fitness of Caulobacter crescentus cells naturally expressing a Vsr-like protein"

### SUPPLEMENTARY DATA AND MATERIAL

**Content:**

Pages 2-12: Supplementary Figures S1 to S11 with legends

Pages 13-17: Supplementary Tables S1 to S3 with captions of Tables S1 to S5

Pages 17-20: Supplementary Material and Methods

Page 20: Supplementary References

**Supplementary Figures with their legends:**

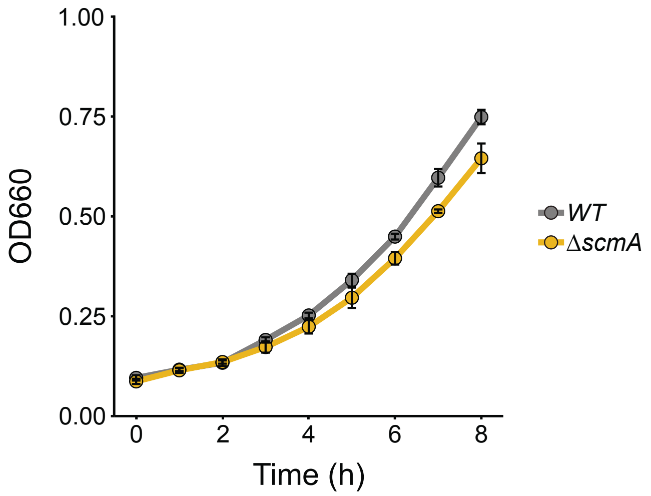


**Figure S1: The growth rates of wild-type and Δ*scmA* *C. crescentus* cells cultivated in minimal medium are globally comparable.** Growth curves of *WT* (JC450) and *ΔscmA* (JC2005) cells cultivated in minimal M2G medium. Cells were first cultivated overnight in PYE medium (reaching stationary phase) and cultures were then diluted 10-fold into M2G medium. Cells were then cultivated until they reached stationary phase, and the cultures were diluted again into fresh M2G medium to reach an $OD_{660nm}$~0.1 at time 0. The $OD_{660nm}$ was then measured every hour for 8 hours. The values plotted in these growth curves correspond to the average measurements from at least three independent biological experiments (error bars = ± SD).

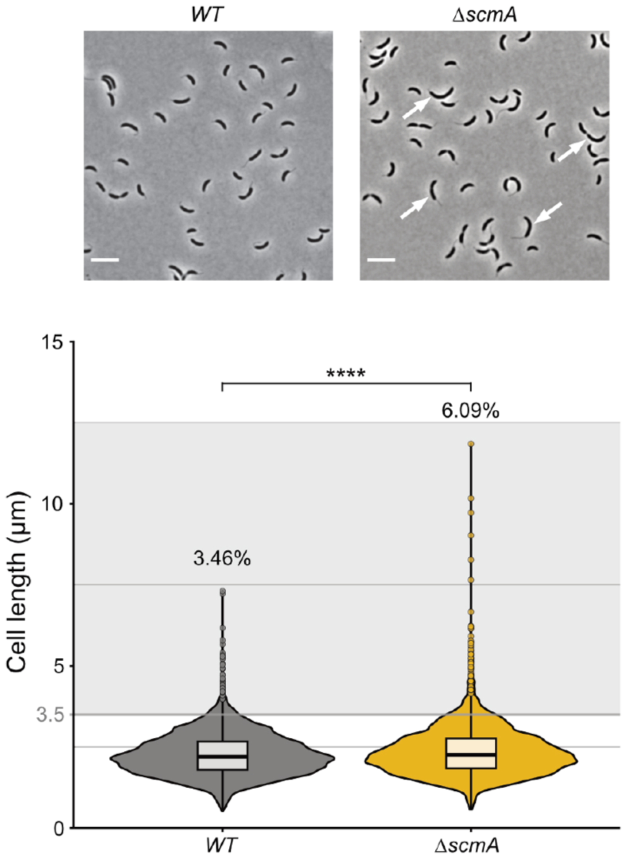


**Figure S2: Δ*scmA* cells are more often significantly more elongated than isogenic wild-type *C. crescentus* cells.** *WT* (JC450) and Δ*scmA* (JC2005) cells were cultivated exponentially in M2G medium until they reached an $OD_{660nm}$~0.4. Cells were then fixed before being imaged by phase-contrast microscopy. Representative images are shown in the upper panel with white arrows pointing at significantly elongated cells. Scale bar = 2 µm. The lower panel corresponds to violin plots showing the cell length distribution in each cell population. The length of minimum 900 cells per biological replicate was measured and the values of three independent biological replicates were used to make these violin plots. Cells greater than 3.5 µm-long (grey zone) were considered as significantly elongated compared to the rest of the population; the percentage of these significantly elongated cells in each population is indicated above each plot. The boxes indicate the interquartile range with the center representing the median. Statistical significance is indicated (****=$p < 0.0001$).

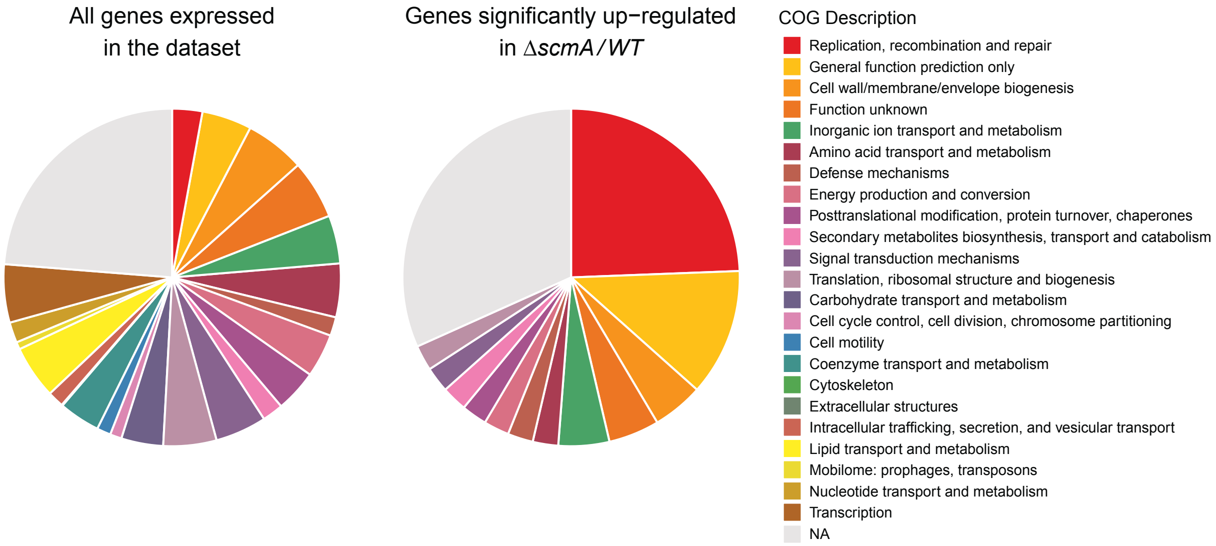


**Figure S3: The COG category "Replication, recombination and repair" is particularly over-represented among the genes that are significantly up-regulated in *ΔscmA* compared to *WT* cells.** Pie charts representing the proportion of genes in each COG category among all the genes that were expressed during these experiments (same experiments as shown in Fig.3) and among the genes that were significantly up-regulated (adjusted *P*-value >0.05 and/or FC<2) in Δ*scmA* (JC2005) cells compared to *WT* (JC450) cells. A distinct color was assigned to each COG category. NA = no COG assigned. COG categories were assigned to *C. crescentus* genes using the *cogclassifier* tool (v1.0.5) from the bioconda. Out of 3'859 UniProtKB entries for the NA1000 strain, 81.58% (3148) protein sequences were classified to a COG functional category.

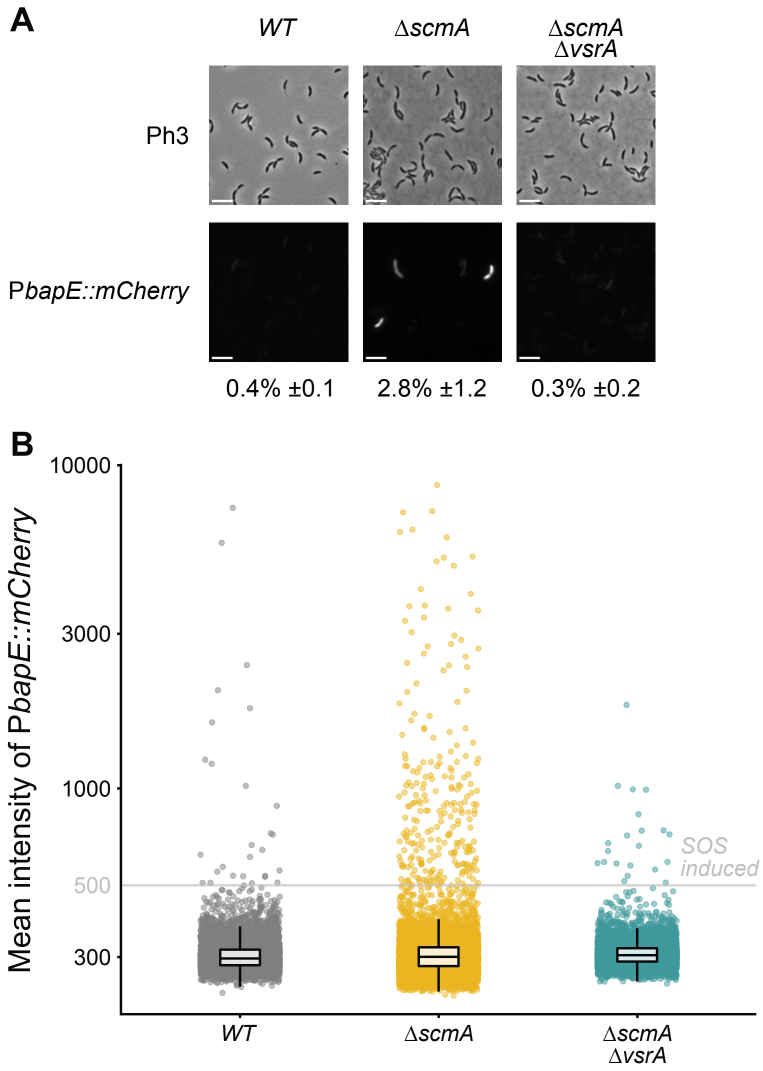


**Figure S4: Single-cell fluorescence microscopy assays showing that a VsrA-dependent SOS response is turned on in a subset of *ΔscmA* cells in clonal populations.** The P*bapE::mCherry* reporter was integrated at the *bapE* locus on the genome of *WT* (giving JC2813), *ΔscmA* (giving JC2814) or *ΔscmA ΔvsrA* (giving JC2815) cells. The resulting strains were cultivated in exponential phase in PYE complex medium. **(A)** Phase contrast (Ph3) and mCherry images of representative cells are shown. Scale bars indicate 2µm. **(B)** Quantification of the cytoplasmic fluorescence intensity (arbitrary units) of cells in populations from (A). The average cytoplasmic fluorescence intensity of minimum 6000 cells of each strain/culture (3 biological replicates of each with minimum 2000 cells/replicate) is shown: the boxes indicate the interquartile range with the center representing the median. Dots above the "SOS" threshold represented by the grey line were used to estimate the percentage of cells in each population displaying an obvious SOS response, as indicated under each microscopy image in panel (A).

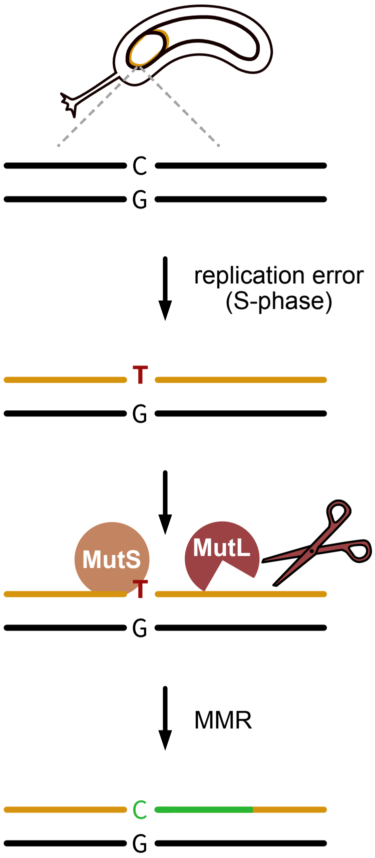


**Figure S5: Model for the DNA mismatch repair (MMR) process in *C. crescentus*** (inspired from (Chai et al. 2021)). During DNA replication (S-phase), the DNA Pol III can accidentally mis-incorporate bases in the newly synthesized DNA strand (represented in orange; a TG mismatch is shown as an example). MutS will recognize these mismatches and recruit/activate the MutL endonuclease to nick the newly replicated DNA strand to initiate the repair of the mismatch. The DNA PolIII then re-synthetizes a patch of that DNA strand (shown in green). How MutL recognizes the newly synthetized DNA strand that it needs to nick during this process remains unclear.

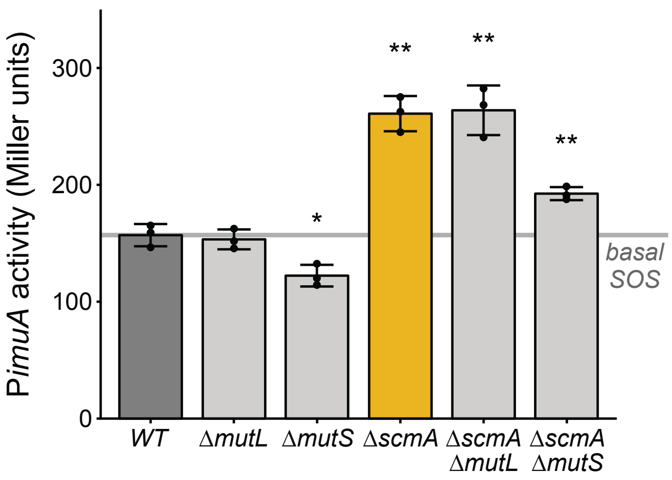


**Figure S6: The SOS response that is turned on in a subset of Δ*scmA* cells does not involve MutL or MutS.** The P*imuA*::*lacZ*290 reporter was introduced into the chromosome of the indicated *C. crescentus* strains and β-galactosidase assays were performed on cells cultivated exponentially in PYE medium (cultures with an $OD_{660nm}$ ~ 0.3). The grey line represents the average basal SOS response found in *WT* cells. The plotted values correspond to the average promoter activities measured from three independent biological replicates with two technical replicates each (error bars = ± SD). Statistical significance is indicated (*=$p < 0.05$; ** =$p < 0.01$).

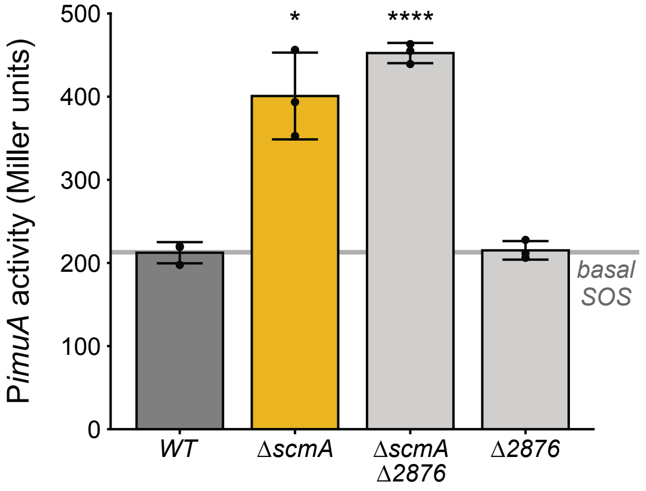


**Figure S7: The CCNA_02876 Vsr-like endonuclease is not involved in the SOS response detected in Δ*scmA* cells.** The P*imuA*::*lacZ*290 reporter was introduced into the chromosome of the indicated *C. crescentus* strains and β-galactosidase assays were performed on cells cultivated exponentially in PYE medium (cultures with an $OD_{660nm}$ ~ 0.4). The grey line represents the average basal SOS response found in *WT* cells. The plotted values correspond to the average promoter activities measured from three independent biological replicates with two technical replicates each (error bars = ± SD). Statistical significance is indicated (*=$p < 0.05$; ****=$p < 0.0001$).

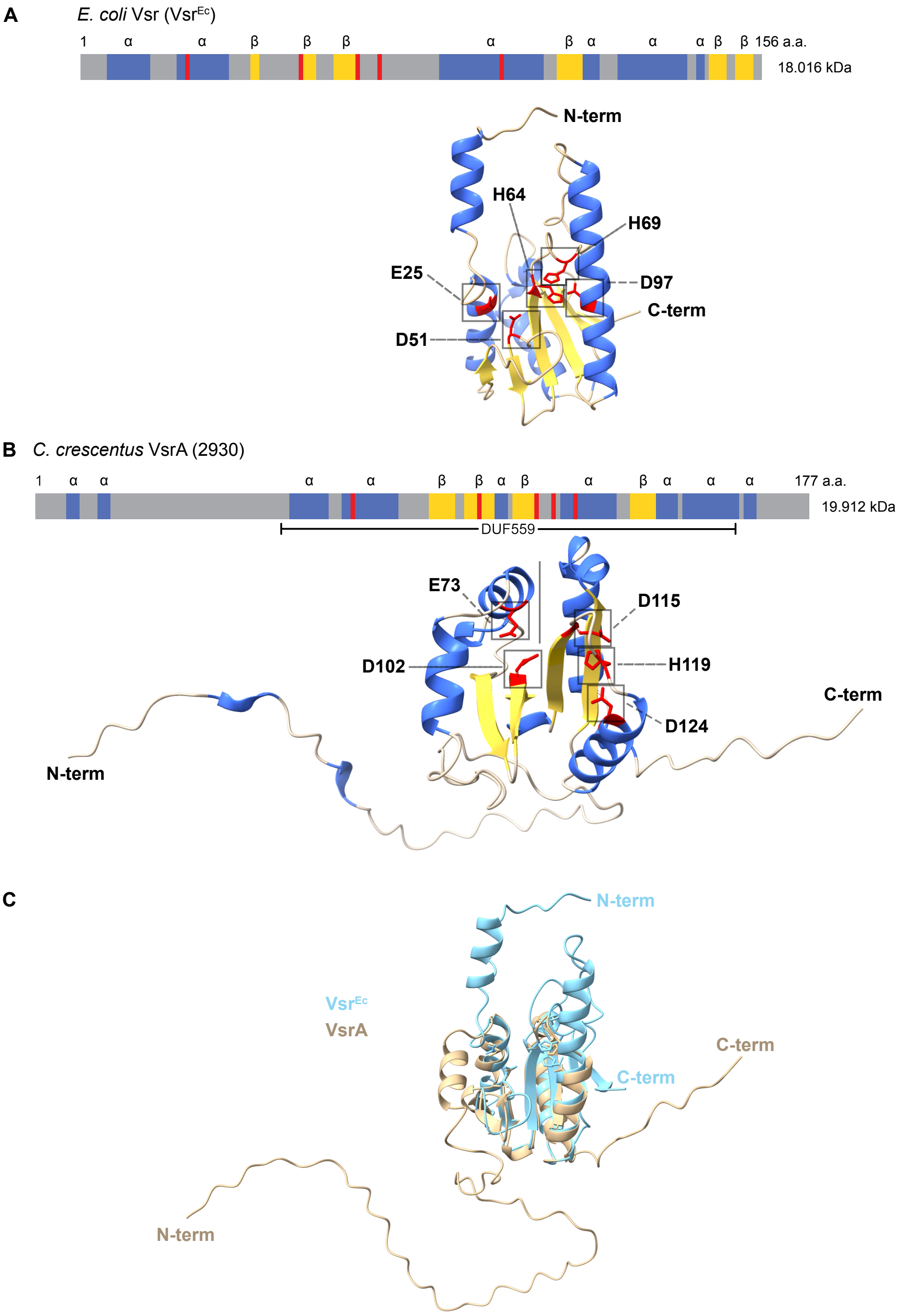


**Figure S8: Comparison of the predicted structures of the *Escherichia coli* Vsr$^{Ec}$ and the *Caulobacter crescentus* VsrA proteins.** Predicted organizations, structures, length in amino-acids and molecular weights of the Vsr$^{Ec}$ protein of the *E. coli* K12 strain **(A)** and of the VsrA (CCNA_02930) protein of the *C. crescentus* NA1000 strain **(B)** are displayed. In panels **A** and **B**, linear and 3D schematics of the predicted structures (Jumper et al. 2021; Varadi and Velankar 2023) are shown. β-strands and α-helices are shown in yellow and blue, respectively. The amino-acid residues that are important for the endonuclease activity of Vsr$^{Ec}$ (E25, D51, H64, H69 and D97) (Tsutakawa et al. 1999a; Tsutakawa et al. 1999b) are highlighted in red. The equivalent residues identified on VsrA (E73, D102, D115, H119 and D124) are also highlighted in red. (**C**) Alignment of the two predicted 3d structures from (A) and (B) with VsrA in brown and $Vsr^{Ec}$ in blue. This superimposition was done using ChimeraX v.1.8 (Meng et al. 2023).

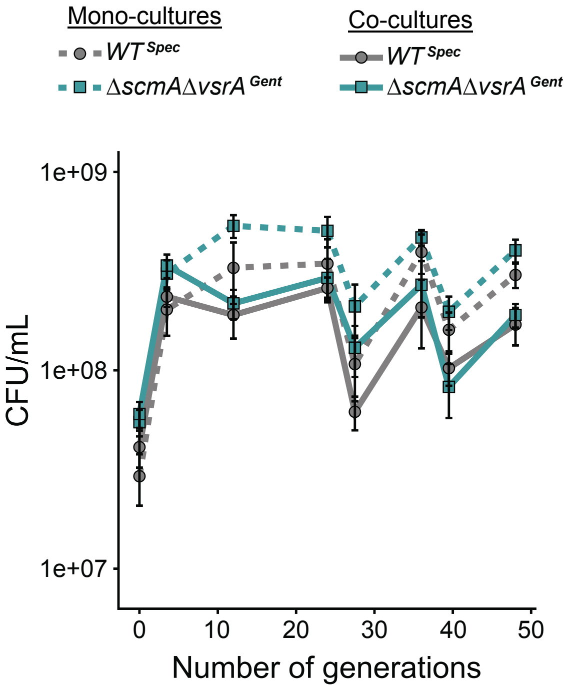


**Figure S9: Wild-type *C. crescentus* cells do not outcompete *ΔscmA ΔvsrA* cells during competition experiments.** Monocultures (dashed lines) and co-cultures (solid lines) of $WT^{Spec}$ (JC2985) and/or Δ*scmA ΔvsrA*$^{Gent}$ (JC3049) cells were cultivated in complex PYE medium for ~48 generations (the generation time of these strains was close to 2 hours under these conditions). Before each regular dilution, spectinomycin-resistant ($WT^{Spec}$) and gentamycin-resistant (Δ*scmA ΔvsrA*$^{Gent}$) colony forming units (CFU) per mL of (co-)cultures were measured on antibiotic-containing PYEA plates. The plotted values correspond to average measurements from four independent biological replicates (error bars = ± SD).

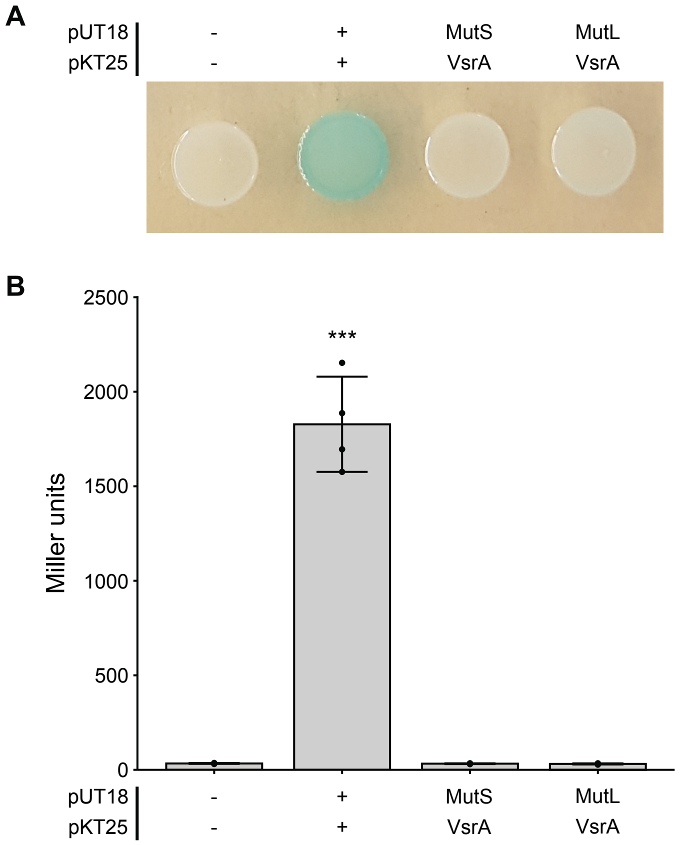


**Figure S10: Bacterial two hybrid assays indicate that VsrA does not interact with the *C. crescentus* MutS or MutL proteins.** MutS and MutL were fused to the T18 fragment of an adenylate cyclase, while VsrA was fused to a T25 fragment to test interactions. Assays were performed using *E. coli* BTH101 cells carrying either the empty pUT18 and pKT25 plasmids, or the pUT18-zip and pKT25-zip plasmids as negative (-) and positive (+) controls, respectively. (**A**) Representative images of bacterial patches on LBA+X-Gal+IPTG plates. (**B**) Results of β-galactosidase assays for BACTH using cells collected from 4 plates as shown in (A). The values are averages of activities measured from four independent biological replicates (error bars = ± SD). Statistical significance is indicated (***=$p < 0.001$).

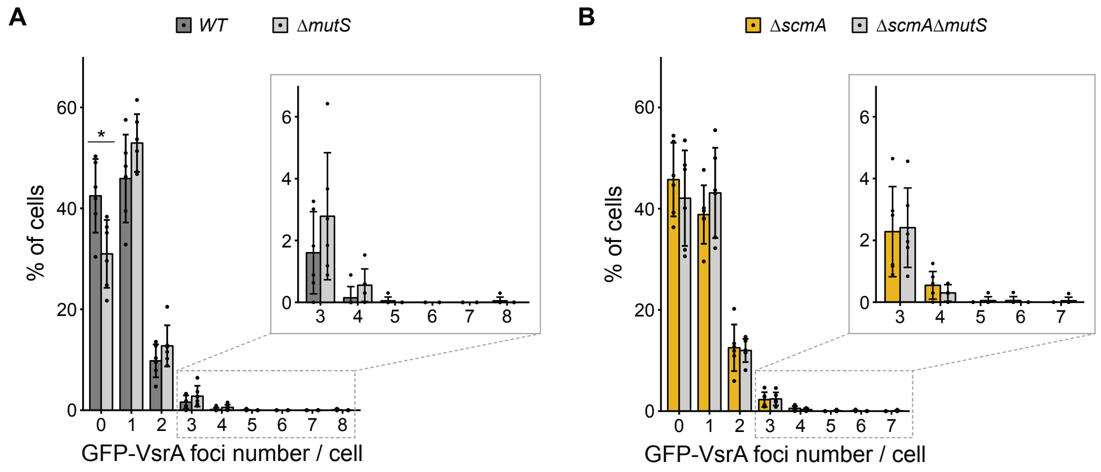


**Figure S11: MutS has no impact on the proportion of cells displaying GFP-VsrA foci in stationary phase.** The native *vsrA* gene was replaced by a *gfp-vsrA* construct in *WT*, Δ*mutS*, Δ*scmA* and Δ*scmA* Δ*mutS* cells, giving the JC2860, JC3072, JC2861 and JC3075 strains, respectively. These strains were then cultivated in PYE medium until they reached stationary phase before being imaged with a fluorescence microscope. Phase contrast and GFP images of cells were acquired and the number of detectable GFP-VsrA foci per cell was analyzed for each cell population. Minimum 300 cells were analyzed for each biological replicate. Panels **(A)** and **(B)** show the percentage of cells displaying a given number of GFP-VsrA foci per cell for each population. The plotted values are averages of at least three independent biological replicates. (error bars = ± SD). Statistically significant differences comparing cells with different genotypes using a student's *t*-test is indicated as follows: * = *P*-value < 0.05.

#### Supplementary Tables:

##### Table S1: Bacterial strains used in this study

| Genotype | Description | Given name | Reference/Origin |
|---|---|---|---|
| *Caulobacter crescentus* | | | |
| NA1000 (WT) | Synchronizable derivative of wild-type strain CB15 (named NA1000 or CB15N) | JC450 | (Evinger and Agabian 1977) |
| NA1000 Δ*scmA* | NA1000 with *scmA* (*CCNA_01085*) deletion | JC2005 | This study |
| NA1000 Δ*vsrA* | NA1000 deleted for *vsrA* (*CCNA_02930*) | JC2540 | This study |
| NA1000 Δ*scmA* Δ*vsrA* | NA1000 Δ*scmA* deleted for *vsrA* | JC2542 | This study |
| NA1000 Δ*mutS* | NA1000 deleted for *mutS* (*CCNA_00012*) | JC1427 | (Chai et al. 2021) |
| NA1000 Δ*scmA* Δ*mutS* | NA1000 Δ*scmA* deleted for *mutS* | JC2987 | This study |
| NA1000 Δ*mutL* | NA1000 deleted for *mutL* (*CCNA_00731*) | JC1426 | (Chai et al. 2021) |
| NA1000 Δ*scmA* Δ*mutL* | NA1000 Δ*scmA* deleted for *mutL* | JC2474 | This study |
| NA1000 Δ*2876* | NA1000 deleted for *CCNA_02876* | JC2539 | This study |
| NA1000 Δ*scmA* Δ*2876* | NA1000 Δ*scmA* deleted for *CCNA_02876* | JC2541 | This study |
| NA1000 P*imuA*::*mCherry* | NA1000 carrying a P*imuA::mCherry* transcriptional fusion at *imuA* native locus | JC3104 | This study |
| NA1000 Δ*scmA* P*imuA*::*mCherry* | NA1000 Δ*scmA* carrying a P*imuA::mCherry* transcriptional fusion at *imuA* native locus | JC3105 | This study |
| NA1000 Δ*vsrA* P*imuA*::*mCherry* | NA1000 Δ*vsrA* carrying a P*imuA::mCherry* transcriptional fusion | JC3107 | This study |
| NA1000 Δ*scmA* Δ*vsrA* P*imuA*::*mCherry* | NA1000 Δ*scmA* Δ*vsrA* carrying a P*imuA::mCherry* transcriptional fusion at *imuA* native locus | JC3109 | This study |
| NA1000 P*bapE*::*mCherry* | NA1000 carrying a P*bapE::mCherry* transcriptional fusion at *bapE* native locus | JC2813 | This study |
| NA1000 Δ*scmA* P*bapE*::*mCherry* | NA1000 Δ*scmA* carrying a P*bapE::mCherry* transcriptional fusion at *bapE* native locus | JC2814 | This study |
| NA1000 Δ*scmA* Δ*vsrA* P*bapE*::*mCherry* | NA1000 Δ*scmA* Δ*vsrA* carrying a P*bapE::mCherry* transcriptional fusion at *bapE* native locus | JC2815 | This study |
| NA1000 *GFP::vsrA* | NA1000 carrying a *GFP::vsrA* translational fusion at *vsrA* native locus | JC2860 | This study |
| NA1000 Δ*scmA* *GFP::vsrA* | NA1000 Δ*scmA* carrying a *GFP::vsrA* translational fusion at *vsrA* native locus | JC2861 | This study |
| NA1000 Δ*mutS* *GFP::vsrA* | NA1000 Δ*mutS* carrying a *GFP::vsrA* translational fusion at *vsrA* native locus | JC3072 | This study |
| NA1000 Δ*scmA* Δ*mutS* *GFP::vsrA* | NA1000 Δ*scmA* Δ*mutS* carrying a *GFP::vsrA* translational fusion at vsrA native locus | JC3075 | This study |

| | | | |
|---|---|---|---|
| NA1000 SpecR | NA1000 with pVGFPC-1 inserted at the *vanA* locus to confer SpecR | JC2985 | This study |
| NA1000 Δ*scmA* GentR | NA1000 Δ*scmA* with pVGFPC-4 inserted at the *vanA* locus to confer GentR | JC2984 | This study |
| NA1000 Δ*scmA* Δ*vsrA* GentR | NA1000 Δ*scmA* Δ*vsrA* with pVGFPC-4 inserted at the *vanA* locus to confer GentR | JC3049 | This study |
| *Escherichia coli* | | | |
| F- *mcrA Δ(mrr-hsdRMS-mcrBC) φ80lacZΔM15 ΔlacX74 nupG recA1 araΔ139 Δ(ara-leu)7697 galE15 galK16 rpsL*($Str^R$) *endA1l*$^-$ | Used for cloning procedures | TOP10 | Invitrogen |
| F *cya*-99 *araD*139 *galE*15 *galK*16 *rpsL*1 ($Str^R$) *hsdR*2 *mcrA*1 *mcrB*1 | Adenylate cyclase-deficient *E. coli* strain used for bacterial adenylate cyclase two-hybrid assays (BACTH) | BTH101 | Euromedex |

#### Table S2: Plasmids constructed and/or used during this study

| **Plasmid name** | **Description** | **Strain name with this plasmid** | **Reference** |
|---|---|---|---|
| pBXMCS-2 | High copy number vector with pBBR1 origin ($Kan^R$) | LS4421 | (Thanbichler et al. 2007) |
| pBX-1motif | pBXMCS-2 plasmid containing a 2200 bp insert flanked by HpaII sites and containing a single YGCCGGCR motif (CGCCGGCG) (KanR) | JC3051 | This study |
| pNPTS138 | Suicide vector carrying the *sacB* gene, with ColEI origin and *oriT* (KanR) used as a tool to delete/mutate genes in *C. crescentus* (cannot replicate in *C. crescentus*) | JC473 | D. Alley, unpublished |
| pNPTS138::Δ*scmA* | pNPTS138 containing *scmA* flanking regions and used to create the *scmA* deletion (KanR) | JC1999 | This study |
| pNPTS138::Δ*vsrA* | pNPTS138 containing *vsrA* flanking regions and used to create the *vsrA* deletion (KanR) | JC2496 | This study |
| pNPTS138::Δ*mutS* | pNPTS138 containing *mutS* flanking regions and used to create the *mutS* deletion (KanR) | JC1282 | (Chai et al. 2021) |
| pNPTS138::Δ*mutL* | pNPTS138 containing *mutL* flanking regions and used to create the *mutL* deletion (KanR) | JC1283 | (Chai et al. 2021) |
| pNPTS138::Δ*2876* | pNPTS138 containing *CCNA_02876* flanking regions and used to create the *CCNA_02876* deletion (KanR) | JC2495 | This study |
| pCHYC-1 | Vector with ColEI origin (not functional in *C. crescentus*) and *oriT* (SpecR, StrepR) used to generate C-terminal mCherry protein fusions and used as | LS4218 | (Thanbichler et al. 2007) |

| | | | |
|---|---|---|---|
| | template to amplify *mCherry* to construct pNPTS138::P*imuA::mCherry* | | |
| pNTPS138::P*imuA:: mCherry* | pNPTS138 derivative used to insert *mCherry* under the control of the native *imuA* promoter in the *C. crescentus* chromosome (KanR) | JC3099 | This study |
| pNTPS138::P*bapE:: mCherry* | pNPTS138 derivative used to insert a*mCherry* under the control of the native *bapE* promoter in the *C. crescentus* chromosome (KanR) | JC2792 | This study |
| pGFPC-1 | Vector with ColEI origin (not functional in *C. crescentus*) and *oriT* used to amplify the *GFP* gene to construct pNPTS138::*GFP::vsrA* (SpecR, StrepR) | LS4213 | (Thanbichler et al. 2007) |
| pNPTS138::*GFP::vsrA* | pNPTS138 derivative used to insert *GFP::vsrA* at the native *vsrA* locus (KanR) | JC2852 | This study |
| pVGFPC-1 | Vector with ColEI origin (not functional in *C. crescentus*) and *oriT* and with the *vanA* promoter used to confer SpecR resistance once integrated at the native *vanA* locus on the *C. crescentus* chromosome (SpecR, StrepR) | LS4229 | (Thanbichler et al. 2007) |
| pVGFPC-4 | Vector with ColEI origin (not functional in *C. crescentus*) and *oriT* and with the *vanA* promoter used to confer GentR resistance once integrated at the native *vanA* locus on the *C. crescentus* chromosome (GentR) | LS4340 | (Thanbichler et al. 2007) |
| p*lacZ*290 | Low copy number vector with RK2 origin and *oriT,* used to create *lacZ* transcriptional fusions ($Tet^R$) | JC452 | (Gober and Shapiro 1992) |
| pP*imuA*::*lacZ*290 | p*lacZ*290 derivative with *imuA* (*CCNA_03319*) promoter inserted upstream of *lacZ* (TetR) | JC1053 | (Galhardo et al. 2005) |
| pUT18 | High copy number vector with ColEI origin, used for BACTH C-terminal T18 fusions (AmpR) | JC2135 | Euromedex |
| pKT25 | Medium copy number vector with p15A origin, used for BACTH C-terminal T25 fusions (KanR) | JC2138 | Euromedex |
| pUT18.zip | pUT18 encoding T18-leucine zipper from yeast GCN4 (AmpR) | JC2137 | Euromedex |
| pKT25-zip | pKT25 encoding T25-leucine zipper from yeast GCN4 (KanR) | JC2139 | Euromedex |
| pUT18::*mutS* | pUT18 encoding MutS-T18 (AmpR) | JC3030 | This study |
| pUT18::*mutL* | pUT18 encoding MutL-T18 (AmpR) | JC3031 | This study |
| pKT25::*vsrA* | pKT25 encoding VsrA-T25 (KanR) | JC3038 | This study |

**Table S3: Oligonucleotides used during this study.** Relevant restriction sites of indicated restriction endonucleases are underlined.

| Primer name | Sequence (5' to 3') | Used for |
|---|---|---|
| NM210 | AATAAGCTTCCGGCCACAAGCCGATCGTC (HindIII) | construction of pBX-1motif |
| NM211 | AAAAGGTACCAGAACGACGCCGGCG (KpnI) | |
| NM212 | ATGGTACCCTCAGGTCGATCAGGCTG (KpnI) | |
| NM213 | ATACTAGTCCGGGAATCCGCAGC (SpeI) | |
| TC79 | TCACTAGTATGTCGATCCTGGTCAACAGCGA (SpeI) | construction of pNPTS138::Δ*scmA* |
| TC80 | GCTGGATCCCACCAATTCCACTCCAAAGTCGA (BamHI) | |
| TC81 | CGAGGATCCGCCTAGATTTCCATGGGTCGGAT (BamHI) | |
| TC82 | CTCGAATTCCCCGCGCTTAAACTGCAATCCG (EcoRI) | |
| NM31 | GGACTAGTGCAGAGAGGACATTGTCCATGC (SpeI) | construction of pNPTS138::Δ*vsrA* |
| NM32 | GGAGCGGCGCAGGCTTCAAGTTAGCAAGCTGCAAAGTG | |
| NM33 | GCCCCCCACTTTGCAGCTTGCTAACTTGAAGCCTGCGCC | |
| NM34 | ATGGATCCTGACCTTCCGGGTGGGAACCTG (BamHI) | |
| NM27 | GGACTAGTTTGCCGAAAAACGCCTCCATC (SpeI) | construction of pNPTS138::Δ*2876* |
| NM28 | CAAATTTTCCGCAGTTAGGGTTATTCGGGTGGCGACATC | |
| NM29 | CGAGAGATGTCGCCACCCGAATAACCCTAACTGCGGAAA | |
| NM30 | CGGGATCCGTTGAAGAGGCCGACTTATTTC (BamHI) | |
| NM218 | GGACTAGTTGGGCTGACAGATCGGAAG (SpeI) | construction of pNPTS138::P*imuA::mCherry* |
| NM219 | TCTCCTCTTTAATTTATCCGAAGCGTCGTCCGG | |
| NM220 | ACGCTTCGGATAAATTAAAGAGGAGAAATACTAGATGGTGAGCAAGGGCGAGGA | |
| NM221 | CTTGGGGGGAGGATTTACTTGTACAGCTCGTCCATG | |
| NM222 | GAGCTGTACAAGTAAATCCTCCCCCCAAGGGGGAGGAG | |
| NM223 | ATAGAATTCGGCGCAATGGCGAGGTCC (EcoRI) | |
| NM114 | GGACTAGTGTTAGCCATCGCGCTTTTCTG (SpeI) | construction of pNPTS138::P*bapE::mCherry* |
| NM115 | TCTCCTCTTTAATTCATCGCGACCAGCAGGCGA | |
| NM116 | CTGGTCGCGATGAATTAAAGAGGAGAAATACTAGATGGTGAGCAAGGGCGAGGA | |
| NM117 | GTGCCTAAGCCCCTTTACTTGTACAGCTCGTCCATG | |
| NM118 | GAGCTGTACAAGTAAAGGGGCTTAGGCACCCTTGG | |
| NM119 | ATAGAATTCGGATATCCTTCACCACCCGCTCAC (EcoRI) | |
| NM169 | GGACTAGTCTGGAGGTTAACGCGTGGCA (SpeI) | construction of pNPTS138::*GFP::vsrA* |
| NM170 | CTTGCTCACCATGCCCGGAAGGGATGGATAGG | |
| NM171 | TATCCATCCCTTCCGGGCATGGTGAGCAAGGGCGAGGA | |
| NM172 | CGGACAACCTCCACCAGCTGCAGCCTTGTACAGCTCGTCCATG | |

| | | |
|---|---|---|
| NM173 | TGCAGCTGGTGGAGGTTGTCCGGGCGATGGGCTC | |
| NM174 | ATAGAATTCGCGCAGGCTTCAAGGATCAG (EcoRI) | |
| NM201 | AAAAAAGGATCCCAACGCCCACGCCACGCC (BamHI) | construction of pUT18::*mutS* |
| NM202 | AAGGTACCCGGGCCGTGAGCAGACCCTTAAG (KpnI) | |
| NM203 | AAAAGGATCCCCCCATCCGCCGCCTGCC (BamHI) | construction of pUT18::*mutL* |
| NM204 | AAAAGGTACCCGCCGCCGCCCGAACAGCTTCTC (KpnI) | |
| NM205 | AAAAGGATCCCTGTCCGGGCGATGGGCTC (BamHI) | construction of pKT25::*vsrA* |
| NM206 | AAAAGGTACCCGTCCGTCGACTGCGTCGACAC (KpnI) | |

**Table S4 (available as supplementary .xls file): RNA-Seq results comparing the transcriptome of Δ*scmA* (JC2005) and WT (JC450) cells.** The complete data and a list of genes that were significantly mis-regulated (adjusted *P* value <0.05 and $log_2FC \geq 1$ or $log_2FC \leq -1$) are included in two distinct sheets. Significantly mis-regulated genes are highlighted in grey. On the sheet containing the list of significant results, genes previously identified as DNA-damage induced genes (Modell et al. 2011; Modell et al. 2014) are highlighted in yellow and the presence or absence of a LexA binding site is stated (SOS response genes). The eventual presence of YGCCGGCR motifs in the 250 bp sequences upstream of the annotated start codon of each gene is also indicated.

**Table S5 (available as supplementary .xls file): RNA-Seq results comparing the transcriptome of Δ*scmA* Δ*vsrA* (JC2542) and Δ*vsrA* (JC2540) cells.** The complete data and a list of genes that were significantly mis-regulated (adjusted *P* value <0.05 and $log_2FC \geq 1$ or $log_2FC \leq -1$) are included in two distinct sheets. Significantly mis-regulated genes are highlighted in grey. The eventual presence of YGCCGGCR motifs in the 250 bp sequences upstream of the annotated start codon of each gene is also indicated.

#### Supplementary Material and Methods

##### Growth conditions

*E. coli* cells were cultivated in/on Luria-Bertani (LB) medium or on LB agar (1.5%) plates at 37°C. *C. crescentus* strains were grown in/on Peptone Yeast Extract (PYE+/-1.5% agar) complex medium or in M2G minimal medium (Ely 1991). When required to maintain plasmids into strains, antibiotics were added into the media at the following concentrations (liquid/plates): kanamycin 30/50 µg/ml, streptomycin 30/30 µg/ml, gentamycin 15/20 µg/ml, oxytetracycline 12/12 mg/ml and ampicillin 50/100 mg/ml for *E. coli*; kanamycin 5/25 µg/ml, spectinomycin 25/100 µg/ml, gentamycin 1/5 µg/ml and oxytetracycline 1/1 µg/ml for *C. crescentus*. During BACTH assays, isopropyl β-d-1-thiogalactopyranoside (IPTG, AppliChem) was added at a final concentration of 0.5 mM and 5-bromo-4-chloro-3-indolyl-β-d-galactopyranoside (X-gal, AppliChem) was added at a final concentration of 40 µg/ml to detect β-galactosidase (LacZ) activity on plates.

##### Molecular cloning procedures

DNA manipulations and molecular cloning procedures were performed according to standard protocols. All constructs were verified by colony PCR and/or Sanger sequencing (Microsynth AG).

**Plasmid constructions**

*Derivative of pBXMCS-2:*

To construct pBX-1motif: a first 878 bp sequence containing a 5'-end HpaII motif and a 3'-end YGCCGGCR motif and a second 1359 bp sequence containing a 3'-end HpaII motif, were both amplified using primer pairs NM210/NM211 or NM212/213 using chromosomal *C. crescentus* NA1000 gDNA as a template. The two fragments were digested with HindIII/KpnI and KpnI/SpeI, respectively, and cloned into the HindIII/SpeI-digested pBXMCS-2 vector by triple ligation.

*Derivatives of pNPTS138:*

-To construct pNPTS138*::ΔscmA*: the ~500 bp sequences upstream and downstream of the *scmA* (*CCNA_01085*) ORF were amplified with primer pairs TC79/TC80 or TC81/TC82 using chromosomal *C. crescentus* NA1000 gDNA as a template. The two fragments were digested with SpeI/BamHI and BamHI/EcoRI, respectively, and cloned into the SpeI/EcoRI-digested pNPTS138 vector by triple ligation.

-To construct pNPTS138*::ΔvsrA*: the ~500 bp sequences upstream and downstream of *vsrA* (*CCNA_02930*) ORF were amplified with primer pairs NM31/NM32 or NM33/NM34 using chromosomal *C. crescentus* NA1000 gDNA as a template. The two fragments contained 33-nucleotide overhangs and were used as templates for overlap extension PCR in a second round of PCR using the primer pair NM31/NM34. The fragment was subsequently digested with SpeI/BamHI and cloned into the SpeI/BamHI-digested pNPTS138 vector by double ligation.

-To construct pNPTS138*::Δ2876*: the ~500 bp sequences upstream and downstream of *CCNA_02876* were amplified with primer pairs NM27/NM28 or NM29/NM30 using chromosomal *C. crescentus* NA1000 gDNA as a template. The two fragments contained a 34-nucleotide overhang and were used as templates for overlap extension PCR in a second round of PCR using primer pair NM27/NM30. The fragment was subsequently digested with SpeI/BamHI and cloned into the SpeI/BamHI-digested pNPTS138 vector by double ligation.

-To construct pNPTS138::P*imuA::mCherry*: the ~500 bp sequences upstream and downstream of the 3'-end of *imuA* were amplified with primer pairs NM218/NM219 or NM222/NM223 using chromosomal *C. crescentus* NA1000 gDNA as a template. The *mCherry* ORF was amplified from pCHYC-1 using the primer pair NM220/NM221 that created a 5'-aaagaggagaaa-3' ribosome binding site located in frame with the *mCherry* ORF. The three fragments containing 26-29-nucleotide overhangs were then used as templates for overlap extension PCR in a second round of PCR using the primer pair NM218/NM223. The fragment was subsequently digested with SpeI/EcoRI and cloned into the SpeI/EcoRI-digested pNPTS138 vector by double ligation.

-To construct pNPTS138::P*bapE::mCherry*: the ~500 bp sequences upstream and downstream of the 3'-end of *bapE* were amplified with primer pairs NM114/NM115 or NM118/NM119 using chromosomal *C. crescentus* NA1000 gDNA as a template. The *mCherry* ORF was amplified from pCHYC-1 using the primer pair NM116/NM117 that created a 5'-aaagaggagaaa-3' ribosome binding site in frame with the *mCherry* ORF. The three fragments containing a 26-29-nucleotide overhangs were used as templates for overlap extension PCR in a second round of PCR using primer pair NM114/NM119. The fragment was subsequently digested with SpeI/EcoRI and cloned into the SpeI/EcoRI-digested pNPTS138 vector by double ligation.

-To construct pNPTS138::*GFP::vsrA*: the ~500 bp sequences upstream and downstream of the 5' end of the *vsrA* ORF were amplified with primer pairs NM169/NM170 and NM173/NM174 using chromosomal *C. crescentus* NA1000 gDNA as a template. The *GFP* gene without its stop codon was amplified from pGFPC-1 using the primer pair NM171/NM172. The three fragments containing 22-30-nucleotide overhangs were used as templates for overlap extension PCR in a second round of PCR using the primer pair NM169/NM174. The fragment was subsequently digested with SpeI/EcoRI and cloned into the SpeI/EcoRI-digested digested pNPTS138 vector by double ligation.

*Derivatives of pUT18:*
-To construct pUT18::*mutS*: the *mutS* ORF was amplified with primer pair NM201/NM202 using chromosomal *C. crescentus* NA1000 gDNA as a template. The fragment was subsequently digested with BamHI/KpnI and cloned into the BamHI/KpnI-digested pUT18 BACTH vector by double ligation.

-To construct pUT18::*mutL*: the *mutL* ORF was amplified with primer pair NM203/NM204 using chromosomal *C. crescentus* NA1000 gDNA as a template. The fragment was subsequently digested with BamHI/KpnI and cloned into the BamHI/KpnI-digested pUT18 BACTH vector by double ligation.

*Derivative of pKT25:*
To construct pKT25::*vsrA*: the *vsrA* ORF was amplified with primer pair NM205/NM206 using chromosomal *C. crescentus* NA1000 gDNA as a template. The fragment was subsequently digested with BamHI/KpnI and cloned into the BamHI/KpnI-digested pKT25 BACTH vector by double ligation.

**Strain constructions**
- Integrative or replicative plasmids were introduced into *C. crescentus* cells by electro-transformation.
- Gene deletions or gene replacements on the *C. crescentus* chromosome were done using pNPTS138 derivatives and a two-step recombination procedure. Briefly, pNPTS138 derivatives were first introduced into the chromosome of *C. crescentus* cells by transformation and a first event of homologous recombination at the targeted locus (selection on PYEA+Km plates). A few colonies were then cultivated over-night in PYE (no Km; plasmid re-excision step via a second homologous recombination event) and subsequently plated on PYEA+Sucrose 3% to select the colonies that lost the *sacB* gene/plasmid. Sucrose-resistant and Km-sensitive colonies were then screened and the genotype of these colonies were differentiated by colony-PCR (one expects to find ~50% of *WT* and ~50% of genetically-modified colonies at that step). This method was used to construct the following strains: JC2005, JC2540, JC2542, JC2987, JC2474, JC2539, JC2541, JC3104, JC3105, JC3107, JC3109, JC2813, JC2814, JC2815, JC2860, JC2861, JC3072 and JC3075.
- To construct the JC2985, JC2984 and JC3049 strains, the pVGFPC-1 and pVGFPC-4 integrative plasmids were inserted at the native *vanA* locus on the *C. crescentus* chromosome by homologous recombination, which was confirmed by PCR (Thanbichler et al. 2007).

**Bacterial adenylate cyclase two-hybrid (BACTH) assays**
*E. coli* BTH101 cells were co-transformed with the pUT18 and pKT25 plasmids or with their derivatives by electroporation. ~50 co-transformants were pooled together and resuspended in 100 µl of LB. 5 µl of this cell suspension was then spotted onto LBA plates containing X-

Gal+IPTG+kanamycin+ampicillin and incubated for 48 hours at 30°C. Blue patches then indicate that the two tested proteins interact.
Quantitative β-galactosidase assays were then performed from these bacterial patches as described in (Jaskolska et al. 2018). Briefly, the bacterial patches were collected and resuspended in 1 ml of sterile phosphate-buffered saline (PBS). The suspensions were then diluted 10-fold and 500 µl of this diluted cell suspension was used to perform a β-galactosidase assay as described in (Miller 1972).

#### Supplementary References:


Chai T, Terrettaz C, Collier J. 2021. Spatial coupling between DNA replication and mismatch repair in *Caulobacter crescentus*. *Nucleic Acids Res* **49**: 3308-3321.

Ely B. 1991. Genetics of Caulobacter crescentus. *Methods Enzymol* **204**: 372-384.

Evinger M, Agabian N. 1977. Envelope-associated nucleoid from *Caulobacter crescentus* stalked and swarmer cells. *J Bacteriol* **132**: 294-301.

Galhardo RS, Rocha RP, Marques MV, Menck CF. 2005. An SOS-regulated operon involved in damage-inducible mutagenesis in *Caulobacter crescentus*. *Nucleic Acids Res* **33**: 2603-2614.

Gober JW, Shapiro L. 1992. A developmentally regulated *Caulobacter* flagellar promoter is activated by 3' enhancer and IHF binding elements. *Mol Biol Cell* **3**: 913-926.

Jaskolska M, Stutzmann S, Stoudmann C, Blokesch M. 2018. QstR-dependent regulation of natural competence and type VI secretion in Vibrio cholerae. *Nucleic Acids Res* **46**: 10619-10634.

Jumper J, Evans R, Pritzel A, Green T, Figurnov M, Ronneberger O, Tunyasuvunakool K, Bates R, Zidek A, Potapenko A et al. 2021. Highly accurate protein structure prediction with AlphaFold. *Nature* **596**: 583-589.

Meng EC, Goddard TD, Pettersen EF, Couch GS, Pearson ZJ, Morris JH, Ferrin TE. 2023. UCSF ChimeraX: Tools for structure building and analysis. *Protein Sci* **32**: e4792.

Miller JH. 1972. *Experiments in Molecular Genetics*. Cold Spring Harbor Laboratory Press, Cold Spring Harbor, NY.

Modell JW, Hopkins AC, Laub MT. 2011. A DNA damage checkpoint in *Caulobacter crescentus* inhibits cell division through a direct interaction with FtsW. *Genes Dev* **25**: 1328-1343.

Modell JW, Kambara TK, Perchuk BS, Laub MT. 2014. A DNA damage-induced, SOS-independent checkpoint regulates cell division in *Caulobacter crescentus*. *PLoS Biol* **12**: e1001977.

Thanbichler M, Iniesta AA, Shapiro L. 2007. A comprehensive set of plasmids for vanillate- and xylose-inducible gene expression in *Caulobacter crescentus*. *Nucleic Acids Res* **35**: e137.

Tsutakawa SE, Jingami H, Morikawa K. 1999a. Recognition of a TG mismatch: the crystal structure of very short patch repair endonuclease in complex with a DNA duplex. *Cell* **99**: 615-623.

Tsutakawa SE, Muto T, Kawate T, Jingami H, Kunishima N, Ariyoshi M, Kohda D, Nakagawa M, Morikawa K. 1999b. Crystallographic and functional studies of very short patch repair endonuclease. *Mol Cell* **3**: 621-628.

Varadi M, Velankar S. 2023. The impact of AlphaFold Protein Structure Database on the fields of life sciences. *Proteomics* **23**: e2200128.
